# A polysaccharide deacetylase enhances bacterial adhesion in high ionic strength environments

**DOI:** 10.1101/2021.04.16.440180

**Authors:** Nelson K. Chepkwony, Yves V. Brun

## Abstract

The adhesion of organisms to surfaces in aquatic environments provides a diversity of benefits such as better access to nutrients or protection from the elements or from predation. Differences in ionic strength, pH, temperature, shear forces, and other environmental factors impact adhesion and organisms have evolved various strategies to optimize their adhesins for their specific environmental conditions. We know essentially nothing about how bacteria evolved their adhesive mechanisms to attach efficiently in environments with different physico-chemical conditions. Many species of Alphaproteobacteria, including members of the order *Caulobacterales*, use a polar adhesin, called holdfast, for surface attachment and subsequent biofilm formation in both freshwater and marine environments. *Hirschia baltica,* a marine member of *Caulobacterales,* produces a holdfast adhesin that tolerates a drastically higher ionic strength than the holdfast produced by its freshwater relative, *Caulobacter crescentus*. In this work, we show that the holdfast polysaccharide deacetylase HfsH plays an important role in adherence in high ionic strength environments. We show that deletion of *hfsH* in *H. baltica* disrupts holdfast binding properties and structure. Increasing expression of HfsH in *C. crescentus* improved holdfast binding in high salinity, whereas lowering HfsH expression in *H. baltica* reduced holdfast binding at high ionic strength. We conclude that HfsH plays a role in modulating holdfast binding at high ionic strength and hypothesize that this modulation occurs through varied deacetylation of holdfast polysaccharides.

## INTRODUCTION

The development of adhesives that perform well on wet surfaces has been a challenge for centuries, yet this problem has been solved multiple times during the evolution of sessile aquatic organisms. These organisms derive multiple benefits from their adhesion to surfaces in aquatic environments such as increased access to nutrients, aerated water, and protection from predation. Aquatic environments can differ in ionic strength, pH, temperature, and shear forces, requiring the evolution of environment-optimized adhesion strategies. For example, mussels, a diverse group of bivalve mollusk species, can attach to surfaces in freshwater, brackish waters, and marine habitats, suggesting a successful evolution of adhesion mechanisms adapted to different ionic environments.^1,2^ Both marine and freshwater mussels produce a fibrous polymeric adhesin structure called the byssus for surface attachment.^1,2^ Mussel byssus-mediated adhesion is one of the best characterized systems for how adhesins interact with wet surfaces in both low and high ionic strength environments.^2,3^

The byssus adhesin contains more than 15 mussel foot proteins (Mfps).^3^ Mfp-3 and Mfp-5 contain several 3,4-dihydroxyphenyl-L-alanine (DOPA) residues, a post-translational hydroxylation of tyrosine that promotes byssal adhesiveness and cohesiveness through hydrogen bonding and oxidative cross-linking.^2–4^ Mfp-3 and Mfp-5 are also rich in lysine residues, which are frequently adjacent to DOPA residues on the protein backbone.^1,3^ Marine mussel Mfps have more DOPA residues than freshwater species (11 - 30% mol in marine *vs* 0.1 - 0.6% mol in freshwater), which is hypothesized to contribute to overcoming the binding inhibition posed by high ionic strength in marine waters.^5–8^ The synergy between the DOPA and lysine residues is thought to improve adhesion in marine environments by displacing hydrated salt ions from the surface and increasing electrostatic interactions.^1–3^ Despite the impressive progress in understanding the mechanistic basis for mussel adhesion in different ionic strength environments, the lack of a genetic system has made it difficult to study the evolution of those mechanisms.

Here, we use genetically tractable, related freshwater and marine species of the order *Caulobacterales* to investigate the evolution of adhesion in these two environments. Most bacteria spend their lives attached to or associated with surfaces. Bacteria attach to surfaces using adhesins, which are mainly composed of polysaccharides, DNA, and/or proteins.^9,10^ The mechanism by which adhesins interact with different surfaces is still unclear, but studies have shown that electrostatic interactions play an important role.^11–14^ In marine environments, bacterial adhesins face high ionic strengths, up to 600 mM, compared to ∼ 0.05 mM in freshwater lakes and ponds.^14,15^ Nevertheless, marine bacteria attach efficiently to surfaces in the ocean, despite shielding of electrostatic forces that contribute to surface adhesion in high ionic strength environments.^16,17^ Therefore, bacteria living in such environments must use adhesins that are adapted to binding at high ionic strength.

Species in the order *Caulobacterales* are found as surface-attached cells growing in diverse environment. Their natural habitat ranges from freshwater and marine aquatic environments to nutrient-rich soil and the rhizosphere.^18,19^ Cells attach permanently to surfaces using a specialized polar adhesin called holdfast.^19,20^ *Caulobacter crescentus*, a freshwater member of the *Caulobacterales,* is a stalked bacterium with a dimorphic cell cycle that fluctuates between a flagellated, motile swarmer and a sessile stalked cell.^19^ A swarmer cell differentiates into a stalked cell by shedding its flagellum and synthesizing holdfast-tipped stalk at the same pole (Fig. 1A). Although the exact composition of the *C. crescentus* holdfast is unknown, it has been shown to contain the monosaccharides *N*-acetylglucosamine (GlcNAc), glucose, 3-*O*-methylglucose, mannose, and xylose,^20,21^ as well as proteins and DNA.^22^ The *C. crescentus* holdfast attaches to surfaces with a strong adhesive force of 70 N/mm^2.23,24^

**Figure 1:**
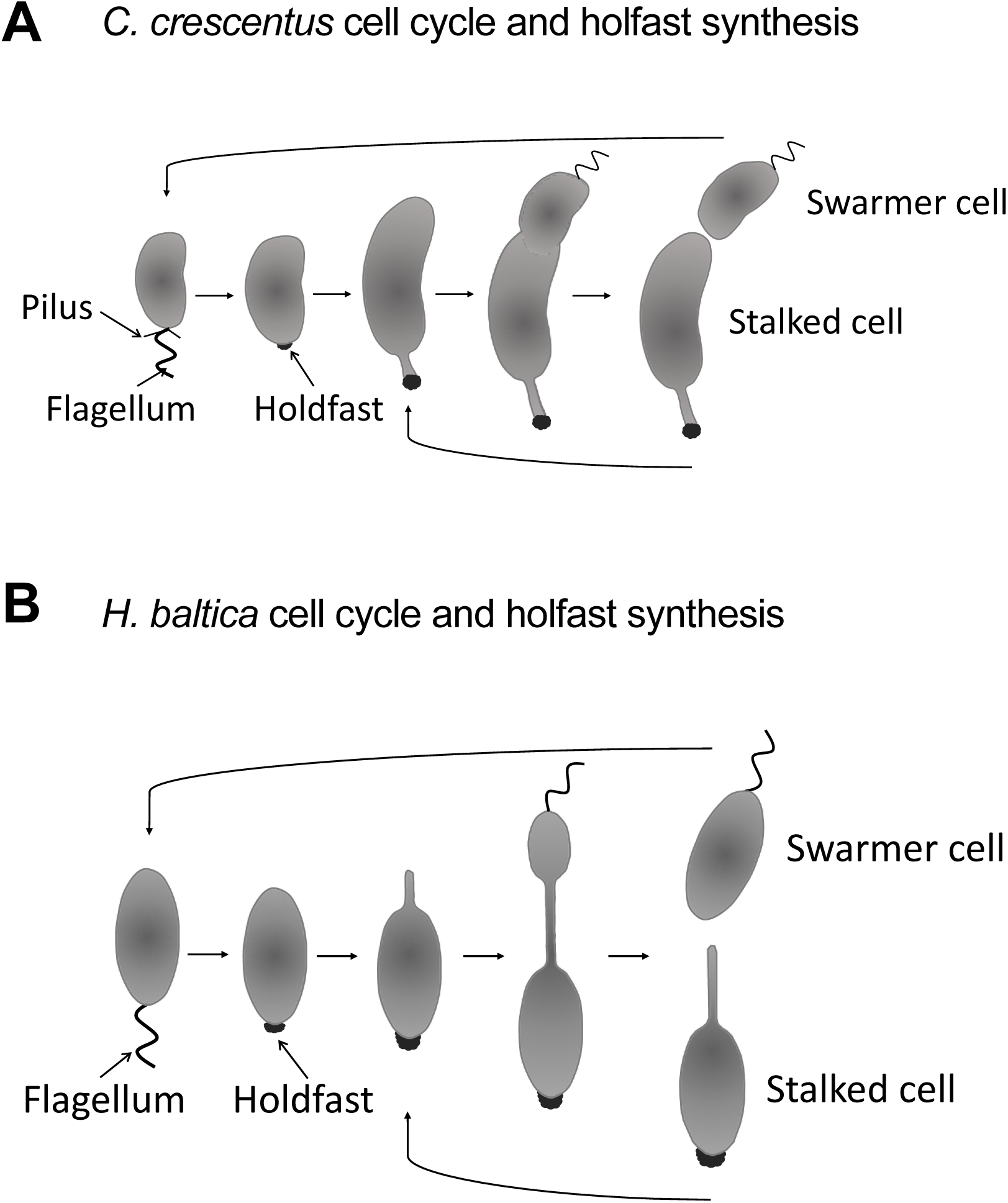
Cell cycle and holdfast synthesis of *C. crescentus* and *H. baltica*. **A**. Diagram of the *C. crescentus* dimorphic cell cycle. A motile swarmer cell differentiates into a stalked cell by shedding the flagellum, retracting the pili, and synthesizing a holdfast-tipped stalk at the same cell pole. *C. crescentus* stalked cells divide asymmetrically to produce a motile swarmer cell and a surface-adherent stalked cell. **B.** Diagram of the *H. baltica* dimorphic cell cycle. A motile swarmer cell differentiates into a stalked cell by shedding its flagellum and synthesizing holdfast at the same cell pole. At the opposite pole, a budding stalk is synthesized that is used to bud a new motile swarmer cell.

*Caulobacterales* use similar genes to synthesize, export, and anchor the holdfast,^9,25^ yet there are substantial differences in holdfast binding properties at high ionic strength. Most studies of holdfast properties have been performed in the freshwater *C. crescentus*,^10^ in which as little as 10 mM NaCl leads to a 50% reduction in binding to glass.^23^ Recently, we studied a marine *Caulobacterales Hirschia baltica,* which produces holdfast at the cell pole and uses the stalk for budding^25^ (Fig. 1B). *H. baltica* produces holdfasts that tolerate a significantly higher ionic strength than *C. crescentus* holdfast, where 600 mM NaCl was required to observe a 50% decrease in binding to glass.^25^ Differences in holdfast tolerance of ionic strength could result from differences in molecular composition or degree or type of modification.

Two holdfast modifying enzymes that have been characterized in the freshwater *C. crescentus* are the putative acetyltransferase, HfsK,^26^ and the putative polysaccharide deacetylase, HfsH.^27^ Deletion of *hfsK* or *hfsH* reduces holdfast cohesiveness (holdfast intramolecular interactions) and adhesiveness (holdfast-surface interactions), which leads to shedding of holdfasts into the medium.^26,27^ This shedding phenotype is similar to that observed in mutants lacking holdfast anchor proteins.^28,29^ Furthermore, overexpression of HfsH in *C. crescentus* increases cell adhesion without increasing the amount of holdfast produced,^27^ implying that not all sugar subunits are deacetylated in wild-type (WT) holdfast. Interestingly, studies on deacetylation of the GlcNAc polymer chitin indicate that removal of acetyl groups leaves the resultant chitosan with an exposed amine group.^30^ The level of deacetylation of chitosan changes its physical and chemical properties by altering electrostatic interactions, acid-base interactions, hydrogen bonds, and hydrophobic interactions with surfaces.^30^ Therefore, we hypothesized that the partial positive charge on the primary amine formed after deacetylation of the holdfast GlcNAc polysaccharide by HfsK and/or HfsH might play a role in improving holdfast binding in high ionic strength environments.

In the present study, we show that the polysaccharide deacetylase HfsH is required for *H. baltica* adhesion and holdfast binding. We demonstrate that holdfast produced by a *H. baltica* Δ*hfsH* mutant is deficient in both cohesive and adhesive properties. *H. baltica* Δ*hfsH* produces a similar quantity of holdfast polysaccharide as the WT, but due to a lack of cohesiveness and adhesiveness, these holdfasts disperse into the medium. Furthermore, we demonstrate that holdfast binding can be modulated by varying the level of expression of HfsH. In *C. crescentus*, overexpression of HfsH increases ionic strength tolerance of holdfasts, while reducing expression of HfsH in *H. baltica* results in reduced ionic strength tolerance. Finally, we show that *H. baltica* HfsH helps to maintain the integrity of the holdfast structure, as holdfasts produced by a *H. baltica* Δ*hfsH* mutant lose their protein and galactose constituents. Collectively our results suggest that modulation of the level of the holdfast polysaccharide deacetylase HfsH is an important adaptation for adherence in high ionic strength environments.

## RESULTS

### The holdfast polysaccharide deacetylase HfsH is required for adhesion and biofilm formation in *H. baltica*

A putative acetyltransferase HfsK, and a polysaccharide deacetylase, HfsH modulate *C. crescentus* holdfast binding properties.^26,27^ In *C. crescentus*, deacetylation of holdfast polysaccharides is important for both the cohesiveness and adhesiveness of holdfast.^27^ In our previous work comparing *H. baltica* and *C. crescentus* holdfasts,^25^ we showed that both species use similar genes to synthesize, export, and anchor holdfast to the cell envelope. We identified *H. baltica* genes that modify holdfast in *C. crescentus*, namely the putative acetyltransferase *hfsK* (*hbal_0069)* and the polysaccharide deacetylase *hfsH* (*hbal_1965*; Fig. 2A). In *C. crescentus,* the *hfsK* gene as well as its paralogs *CC_2277* and *CC_1244* are found outside the core *hfs* locus (Fig. 2A). Similar to *C. crescentus,* the *H. baltica hfsK* gene and its paralogs *hbal_1607* and *hbal_1184* are also found outside the *hfs* locus (Fig. 2A). BLAST analysis did not identify any additional *hfsK* paralogs in *H. baltica*.

**Figure 2:**
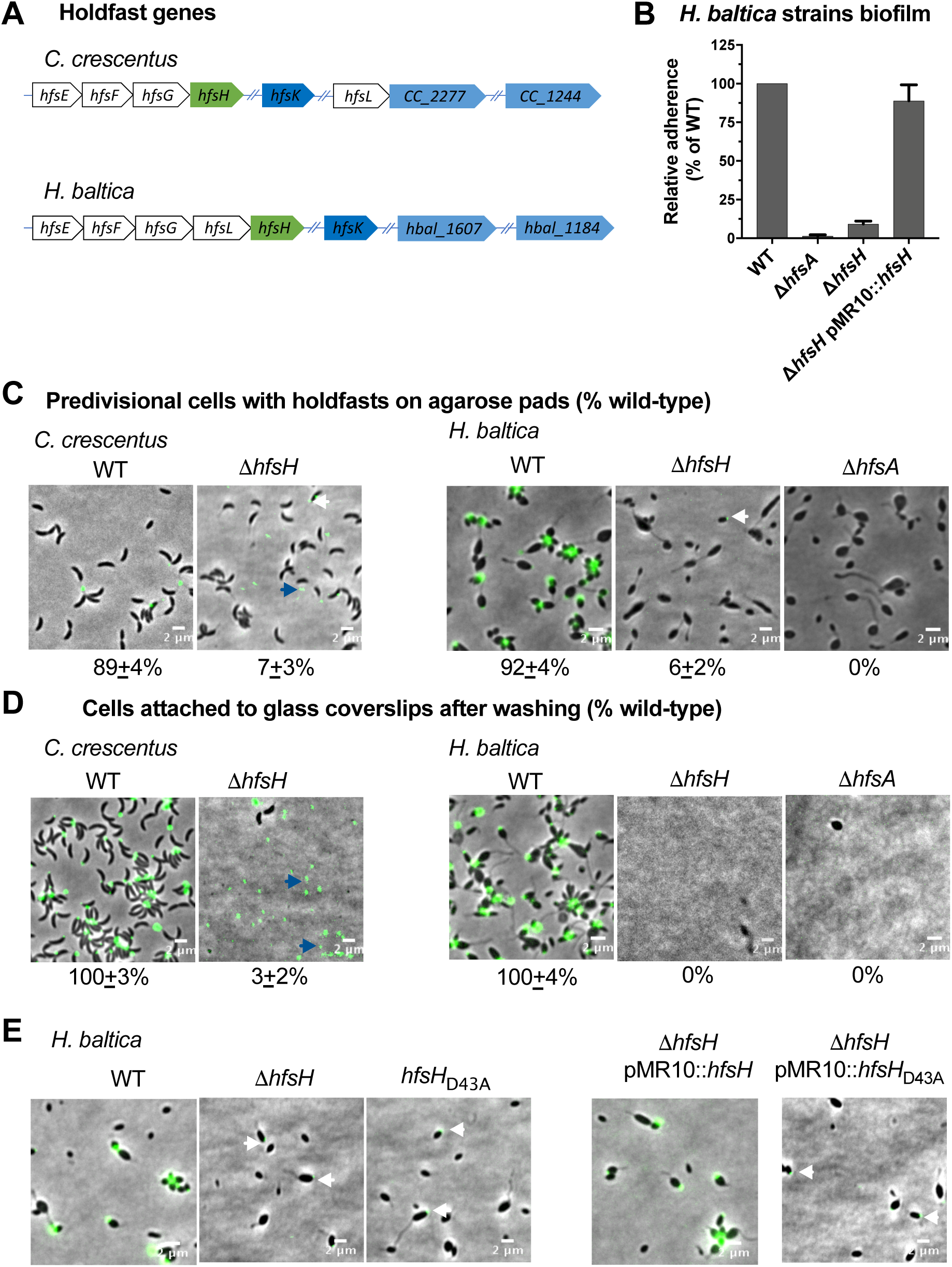
The role of HfsH and HfsK in *H. baltica* holdfast biogenesis. **A.** Genomic organization of holdfast synthesis (*hfs)* genes in *C. crescentus* and *H. baltica.* Genes were identified using reciprocal best hit analysis on *C. crescentus* and *H. baltica* genomes. In both *C. crescentus* and *H. baltica* genomes*, hfsH* is found in the *hfs* locus while *hfsK* and its paralogs are found outside the *hfs* locus. Color coding corresponds to homologs and paralogs. Hash marks indicate genes that are found in a different location in the genome. **B.** Quantification of biofilm formation by the crystal violet assay after incubation for 12 h, expressed as a mean percent of WT crystal violet staining. Holdfast null strain *ΔhfsA* was used as a negative control. Error is expressed as the standard error of the mean of three independent biological replicates with four technical replicates each. **C.** Representative images showing merged phase and fluorescence channels of the indicated *C. crescentus* and *H. baltica* strains on agarose pads. Holdfast is labeled with AF488-WGA (green). White arrows indicate holdfasts attached to the Δ*hfsH* cells, and blue arrows indicate holdfast shed into the medium. Exponential planktonic cultures were used to quantify the percentage of predivisional cells with holdfast. Data are expressed as the mean of three independent biological replicates with four technical replicates each. Error bars represent the standard error of the mean. A total of 3,000 cells were quantified per replicate using MicrobeJ. **D.** Representative images showing merged phase and fluorescence channels of *C. crescentus* and *H. baltica* strains bound to a glass coverslip. Exponential cultures were incubated on the glass slides for 1 h, washed to remove unbound cells, and holdfast were labelled with AF488-WGA (green). Blue arrows indicate surface-bound holdfasts shed by *hfsH* mutants. The data showing quantification of cells bound to the glass coverslip are the mean of two biological replicates with five technical replicates each. Error is expressed as the standard error of the mean using MicrobeJ. **E.** Representative images showing merged phase and fluorescence channels of *H. baltica* strains with holdfast polysaccharides labeled with AF488-WGA (green) on agarose pads. A point mutation was introduced at a key substrate binding residue in *H. baltica* HfsH, resulting in an amino acid change from aspartic acid to alanine at position 43 (D43A). White arrows indicate faint AF488-WGA holdfast labeling on mutant cells

We showed that HfsK is not required for holdfast synthesis, anchoring to the cell envelope, and biofilm formation (see supplementary materials and Fig. S1). Next, we examined the role of the polysaccharide deacetylase HfsH (Hbal_1965) in *H. baltica* biofilm formation by generating an in-frame deletion mutant of *hfsH*. *H. baltica ΔhfsH* was deficient in biofilm formation, similarly to a holdfast null strain *ΔhfsA* (Fig. 2B), and this phenotype could be restored by complementation in *trans* by a replicating plasmid encoding a copy of the *H. baltica hfsH* gene under the control of its native promoter (Fig. 2B). These results show that the polysaccharide deacetylase HfsH plays a significant role in biofilm formation in *H. baltica*.

As holdfast is required for biofilm formation in *C. crescentus* and *H. baltica,*^20,25,31^ we probed for the presence of holdfasts using fluorescent Alexa Fluor 488 (AF488) conjugated <u>w</u>heat <u>g</u>erm <u>a</u>gglutinin (WGA) lectin that specifically binds to the GlcNAc component of the holdfast polysaccharide.^20^ In exponentially growing planktonic cultures, *C. crescentus* WT cells produced holdfasts that bound AF488-WGA and formed cell-cell aggregates mediated by holdfasts, called rosettes (Fig. 2C, left panel). *C. crescentus ΔhfsH* produced holdfasts which associated with the cell or were shed into the medium, as previously shown.^27^ In planktonic culture, *H. baltica* WT formed rosettes and produced holdfasts that bound AF488-WGA and are associated to the cells (Fig. 2C, right panel). *H. baltica* Δ*hfsH* did not form rosettes, and only 6 % of the cells showed some AF488-WGA staining, while many had no AF488-WGA staining similarly to holdfast null mutant Δ*hfsA*, suggesting that no holdfast was associated with these cells (Fig. 2C, right panel, white arrows). Furthermore, we did not observe shed holdfast in the medium from *H. baltica ΔhfsH* (Fig. 2C, right panels), which contrasts with the *C. crescentus ΔhfsH* phenotype (Fig. 2C, left panels, white and blue arrows). Holdfast and rosette formation could be restored in the Δ*hfsH* mutants by complementing in *trans* with a replicating plasmid encoding a copy of the *hfsH* gene (Fig. S2B). We observed that *C. crescentus ΔhfsH* produced small holdfasts which bound to coverslips (Fig. 2D, left panels, blue arrows) but failed to anchor the cells (3% of WT, Fig. 2D), as previously shown.^27^ In contrast, *H. baltica* Δ*hfsH* cells did not bind to the coverslip and we could not detect AF488-WGA labeling on the coverslip surface similarly to a holdfast null mutant Δ*hfsA* (Fig. 2D, right panel), suggesting that holdfasts failed to attach to the glass surface.

HfsH is a predicted carbohydrate esterase family 4 (CE4) enzyme.^27^ The CE4 family of polysaccharide deacetylases have five catalytic motifs for substrate and co-factor binding, as well as those that participate directly in the catalytic mechanism,^32^ which are all present in *C. crescentus* HfsH,^27^ and in *H. baltica* HfsH (Fig. S2D). In order to test if *H. baltica* HfsH is a holdfast polysaccharide deacetylase, we engineered a point mutation in a key substrate-binding residue, resulting in an amino acid change from aspartic acid to alanine at position 43 (D43A; Fig. S2D, asterisk). We monitored for the presence of holdfast using fluorescence microscopy with AF488-WGA. Introduction of D43A in the *H. baltica hfsH* gene phenocopied the *hfsH* deletion (Fig. 2E, white arrows). We complemented the *ΔhfsH* mutant with a WT copy of *hfsH,* or with the point mutant *hfsH_D43A_*, expressed under the native promoter on a low copy replicating plasmid (pMR10). Although WT *hfsH* and *hfsH_D43A_* were expressed similarly (Fig. S2E), complementation with the WT allele restored AF488-WGA holdfast labeling in the *ΔhfsH* mutant background, while the point mutant *hfsH_D43A_* did not (Fig. 2E). These results confirm that *H. baltica* HfsH is involved in holdfast biogenesis and that D43 is important for its activity, similarly to *C. crescentus* HfsH.^27^

The above results indicate that HfsH plays an important role in *H. baltica* holdfast properties, including their anchoring to the cell envelope. We hypothesized that (1) *H. baltica* Δ*hfsH* produces a small amount of holdfast polysaccharide that is insufficient to anchor the cell to the surface, similarly to the under-expression of a glycosyltransferase HfsL,^25^ or (2) holdfasts produced by *H. baltica* Δ*hfsH* are deficient in adhesion and/or cohesion and are thus dispersed into the medium.

### Holdfast produced by *H. baltica ΔhfsH* forms thread-like fibers that diffuse into the medium

The deacetylase mutant *H. baltica* Δ*hfsH* showed a more severe holdfast attachment deficiency compared to *C. crescentus* Δ*hfsH* (Fig. 2C-D). Although we did not observe any holdfast binding to glass coverslips for the *H. baltica* Δ*hfsH* mutant, cells grown planktonically had a faint AF488-WGA labeling (Fig. 2C). These results prompted us to perform time-lapse microscopy to better understand production of holdfast by *H. baltica ΔhfsH*. To visualize holdfast production, we spotted exponential-phase cells on top of a soft agarose pad containing AF488-WGA and collected images every 5 min for 12 h. We observed that *H. baltica* WT produced holdfasts that were labelled with AF488-WGA (Fig. 3A, upper panels) and that the *ΔhfsH* mutant initially produced holdfasts similarly to WT (Fig. 3A, lower panels). However, the holdfasts produced by *H. baltica ΔhfsH* appeared more diffuse compared to WT over time (Fig. 3A and Movie S1A-B). These results show that *H. baltica ΔhfsH* produces holdfast material, indicating that HfsH is not essential for holdfast synthesis.

**Figure 3:**
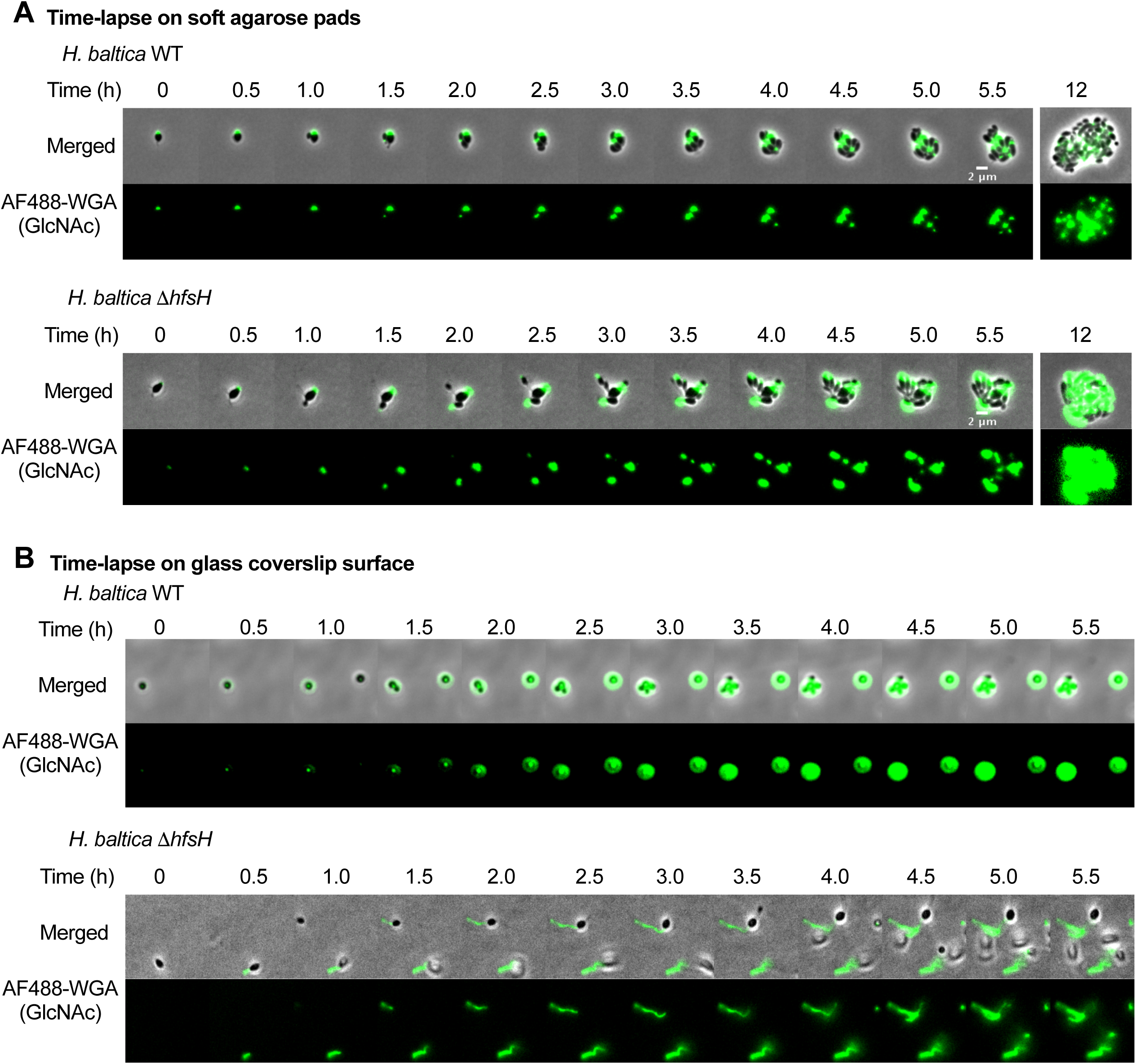
*H. baltica hfsH* mutant holdfasts forms thread-like fibers that diffuse into the medium. **A.** Time-lapse montages of *H. baltica* WT and *H. baltica* Δ*hfsH* on soft agarose pads. Exponential cultures were placed on soft agarose pads containing holdfast-specific AF488-WGA (green) and covered with a glass coverslip. Images were collected every 5 min for 12 h. **B.** Time-lapse montages of *H. baltica* WT and *H. baltica* Δ*hfsH* in microfluidic channels. Exponential cultures with holdfast-specific AF488-WGA (green) were injected into the microfluidic chambers and flow was turned off. Images were collected every 20 sec for 5.5 h.

In order to test how holdfast produced by *H. baltica ΔhfsH* interacts with a glass surface, we performed time-lapse microscopy using a microfluidic device with a low flow rate (1.4 µl/min). We injected exponential-phase cells mixed with AF488-WGA into the microfluidic chamber, turned off the flow, and imaged the cells every 20 sec for 5.5 h. In the microfluidic chamber, we observed that *H. baltica* WT cells arrived at the glass surface and produced holdfasts, allowing them to remain bound to the surface (Fig. 3B, upper panels and Movie S1C). In contrast*, H. baltica ΔhfsH* produced holdfasts that did not remain cohesive on the glass surface and instead formed thread-like fibers (Fig. 3B, lower panel and Movie S1D). These results indicate that HfsH in *H. baltica* is not required for holdfast synthesis, but is essential for maintenance of holdfast cohesive properties.

### HfsH expression correlates with the level of biofilm formation

In order to understand the role of HfsH in *H. baltica,* we investigated whether varying the level of its expression in *H. baltica* affects holdfast cohesive and adhesive properties. We used a copper inducible promoter to tightly control the level of *hfsH* expression^25^ (Fig. 4A). The ability to tightly regulate the expression levels of HfsH under the control of P*_Cu_* was validated by western blot analysis (Fig. S3). We quantified the amount of biofilm formed after 4 h of *hfsH* induction with different concentrations of CuSO_4_ as an inducer of *hfsH* expression. Our results showed that the level of *hfsH* expression in *H. baltica* correlated logarithmically with the relative level of biofilm formed (Fig. 4B). At the highest level of induction at 250 µM CuSO_4_, we observed full restoration of biofilm formation to WT levels (Fig. 4B).

**Figure 4:**
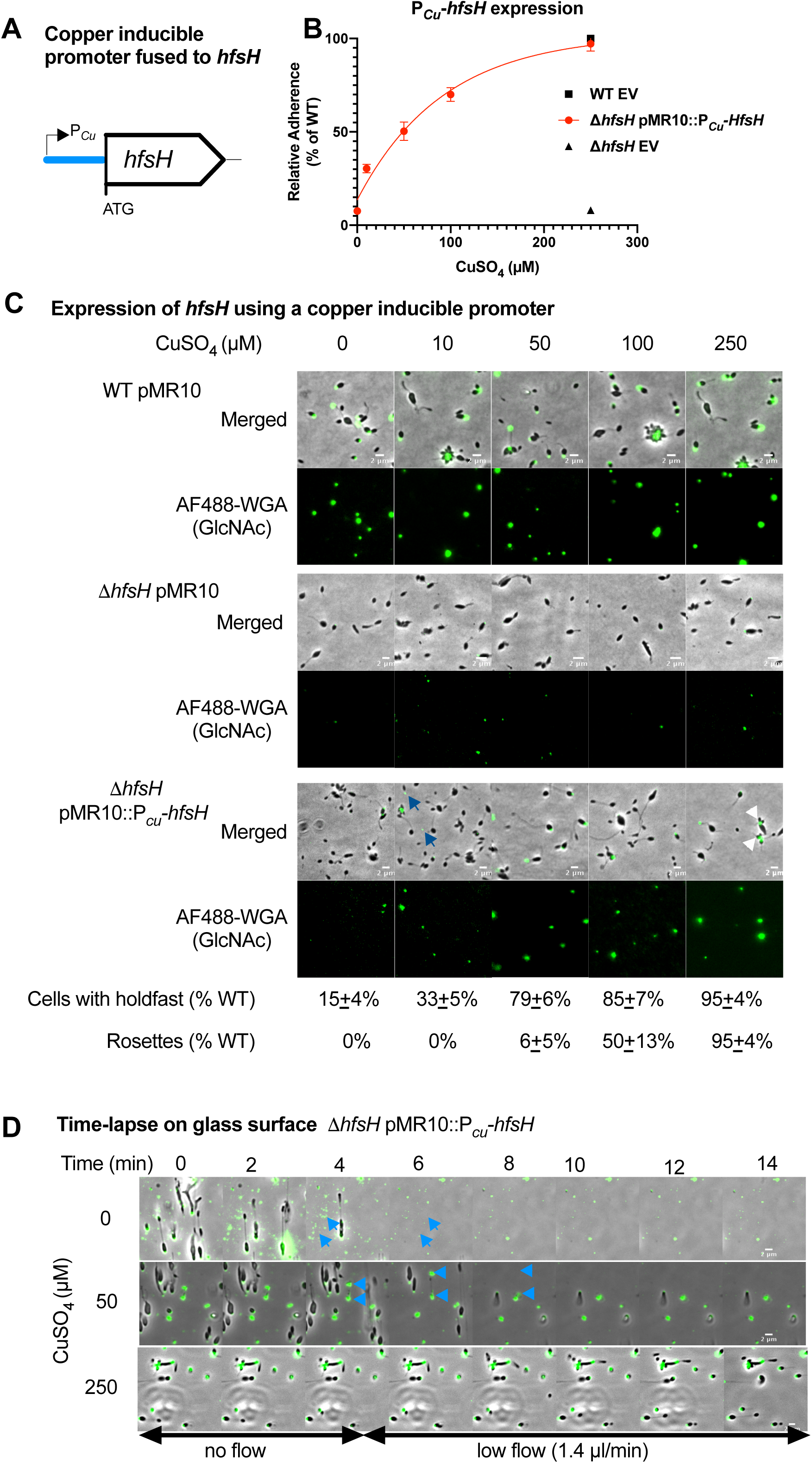
HfsH expression correlates to the level of biofilm formation. **A.** Schematic representation of *hfsH* under the control of a copper inducible promoter. 500 bp upstream of the *copA* open reading frame corresponding to the promoter region, P*_Cu_*, were fused to *hfsH* from *H. baltica* and assembled into the plasmid pMR10. **B.** A logarithmic plot showing quantification of adhesion of *H. baltica* strains induced with 0 – 250 µM of CuSO_4_ for 4 h by the crystal violet assay. Data is expressed as a mean percent of WT crystal violet staining from three independent biological replicates with four technical replicates. Error is expressed as the standard error of the mean. EV is empty vector (pMR10). **C.** Representative images of *H. baltica* WT*, H. baltica* Δ*hfsH*, and *H. baltica* Δ*hfsH* complemented with pMR10::P*_Cu_-hfsH*. Holdfasts were labeled with AF488-WGA (green). Exponential cultures were induced for 2 h with 0 – 250 µM of CuSO_4_. Blue arrows indicate shed holdfast at low levels of induction (10 µM CuSO_4_), and white arrowheads indicate rosettes formed at high levels of HfsH induction (250 µM CuSO_4_). **D.** Time-lapse montages of *H. baltica* Δ*hfsH* pMR10::P*_Cu_-hfsH* in microfluidic channels with holdfast labeled with AF488-WGA (green). Exponential cultures were induced with 0 µM, 50 µM, or 250 µM CuSO_4_, mixed with AF488-WGA, injected into the microfluidic chambers, and allowed to bind for 30 min. Thereafter, the flow rate was adjusted to 1.4 ul/min. Images were collected every 20 sec for 1 h. Blue arrows indicate shed holdfasts.

We next labelled holdfasts with AF488-WGA to analyze the effect of varying HfsH expression on *H. baltica* holdfast production. We induced the expression of HfsH for 2 h in *H. baltica* Δ*hfsH* P*_Cu_-hfsH* using 0 to 250 µM CuSO_4_, and visualized holdfasts of planktonic cells with AF488-WGA by fluorescence microscopy. Addition of CuSO_4_ to *H. baltica* WT and *H. baltica* Δ*hfsH* with empty vector controls had no effect on cell anchoring or holdfast surface adhesion (Fig. 4C, upper and middle panels). In the *H. baltica* Δ*hfsH* mutant complemented with P*_Cu_-hfsH*, we observed a small area of AF488-WGA staining on cells and shed holdfasts in the medium at the lowest level of *hfsH* induction (10 µM CuSO_4_; Fig. 4C, lower panels, blue arrow). As we increased the level of *hfsH* expression, we observed increasing levels of AF488-WGA labeling co-localized with cells (15% at 0 µM CuSO_4_ to 95% at 250 µM CuSO_4_ compared to WT, Fig. 4C, lower panels). At intermediate levels of *hfsH* expression (50 µM CuSO_4_), we observed cells with labeled holdfasts but fewer rosettes compared to full induction at 250 µM CuSO_4_ (6% at 50 µM CuSO_4_ to 95% at 250 µM CuSO_4_, Fig. 4C, lower panels). At the highest level of induction (250 µM CuSO_4_), we observed cells in rosettes and holdfast formation similar to WT (Fig. 4C, lower panels, white arrow), suggesting that holdfast properties were fully restored at this level of HfsH expression.

To test if HfsH expression levels correlate with holdfast cohesiveness and adhesiveness, we performed time-lapse microscopy on *H. baltica* Δ*hfsH* complemented with pMR10::P*_Cu_-hfsH* grown in a microfluidic device. We induced the expression of HfsH for 2 h in liquid cultures, injected exponential-phase induced cells mixed with AF488-WGA into the microfluidic chamber, and turned off the flow for 30 min to allow for binding to the chamber surface. We then adjusted the flow rate to 1.4 µl/min and imaged the cells every 20 sec for 30 min. *H. baltica* Δ*hfsH* pMR10::P*_Cu_-hfsH* grown without CuSO_4_ produced small holdfasts that adhered to the chamber, but were unable to anchor the cells to the surface once flow was turned back on (Fig. 4D, upper panels and Movie S2A). Furthermore, holdfast material that was initially attached to the surface was subsequently washed away upon initiation of fluid flow, suggesting an adhesion defect (Fig. 4D, upper panels, blue arrows). At 50 µM CuSO_4_, we observed a partial restoration of holdfast adhesiveness and cohesiveness as cells were able to stay attached to the surface for longer after re-initiation of the flow, however holdfast adhesiveness was still impaired as shed holdfasts could be easily washed off the surface (Fig. 4D, middle panels, blue arrows and Movie S2B). At 250 µM CuSO_4_, we observed full restoration of holdfast adherence, cell anchoring, and the formation of rosettes (Fig. 4D, lower panels and Movie S2C). These results suggest that at lower levels of HfsH expression holdfast binding properties are only partially restored, while at higher levels of expression holdfast adhesiveness and cohesiveness are fully restored.

### Overexpression of HfsH increases biofilm formation in *C. crescentus* but not in *H. baltica*

*C. crescentus* holdfast binding properties can be increased by overexpressing HfsH,^27^ implying that holdfast polysaccharides are partially deacetylated. *C. crescentus* Δ*hfsH* and *H. baltica* Δ*hfsH* showed important differences in their holdfast structure and binding properties (Fig. 2C-D). Therefore, we hypothesized that there could be differences in the level of holdfast polysaccharide deacetylation among the *Caulobacterales* species. Unfortunately, because it is produced in such small quantities, it is not currently possible to directly determine the level of deacetylation in holdfast. As an alternative, we examined the effect of varying the level of HfsH expression on holdfast adhesive properties. We first tested whether heterologous expression of HfsH in each species would alter holdfast properties. We made two types of cross-complementation constructs for each respective host species: (1) native levels of *hfsH* expression driven by the *hfsH* promoter (P*hfsH_CC_* for *C. crescentus ΔhfsH* and P*hfsH_HB_* for *H. baltica ΔhfsH*), and (2) *hfsH* overexpression driven by the inducible xylose promoter (P*xyl_CC_* and P*xyl_HB_*; Fig. 5A). We analyzed the level of HfsH expression by western blot analysis and found that both HfsH_HB_ and HfsH_CC_ were equally expressed from these promoters (Fig. S4).

**Figure 5:**
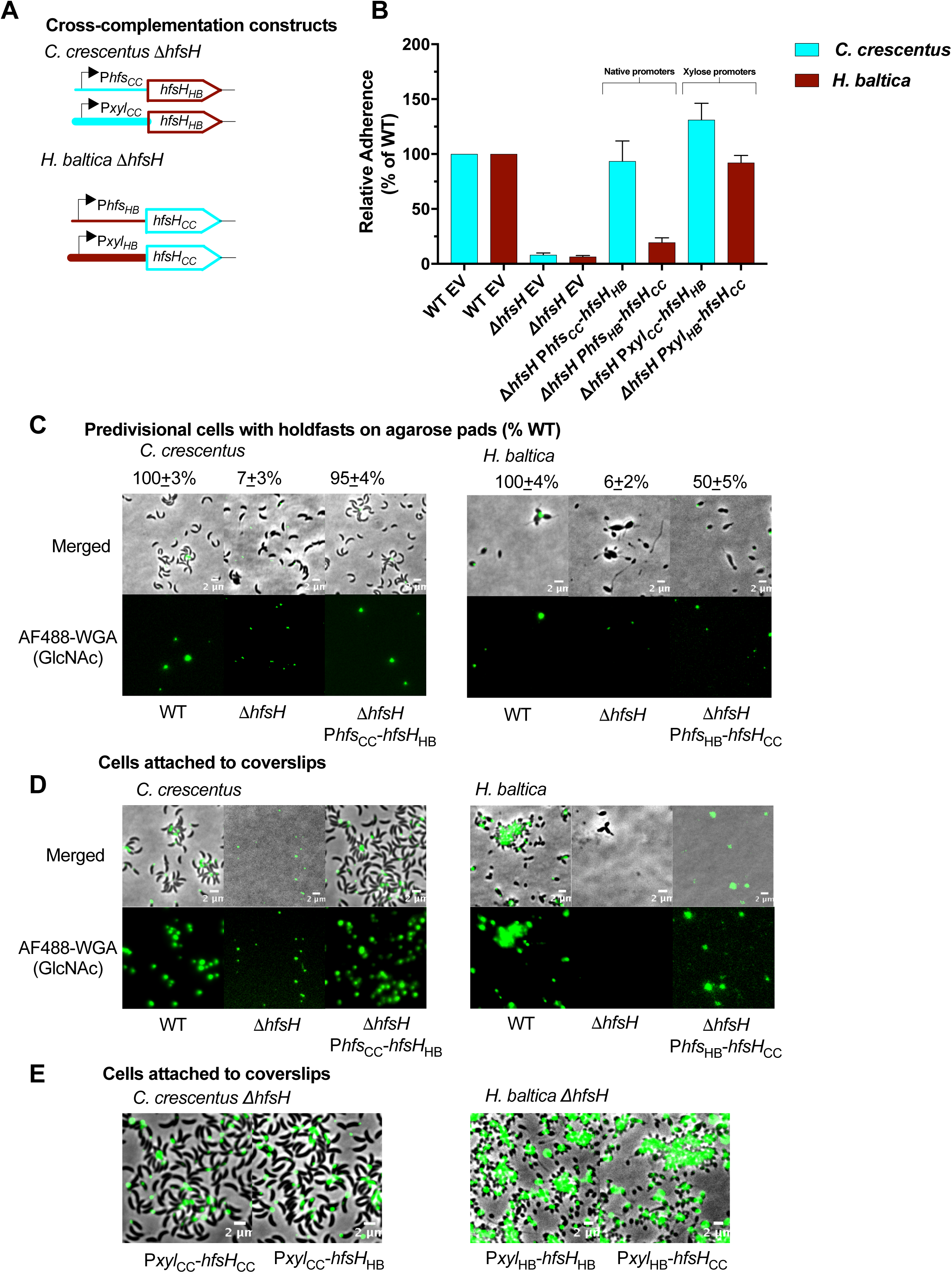
Overexpression of HfsH increases biofilm formation in *C. crescentus* but not *H. baltica*. **A.** Schematic representations of cross-complementation constructs of the *hfsH* gene from *C. crescentus* (*hfsH_CC_*) and *H. baltica* (*hfsH_HB_*) under native holdfast synthesis (P*hfs*) or xylose inducible (P*xyl*) promoters. Native promoters were fused to foreign *hfsH* genes (P*hfs_CC_* and P*xyl_CC_* from *C. crescentus*, or P*hfs_HB_* and P*xyl_HB_* from *H. baltica)* and assembled into the pMR10 plasmid. **B.** Quantification of short-term adhesion (12 h) by the crystal violet assay. Data is expressed as a mean percent of WT crystal violet staining from three biological replicates with four technical replicates each. Error is expressed as the standard error of the mean. **C.** Representative images showing merged phase and fluorescence channels of *C. crescentus* and *H. baltica* strains. Holdfasts are labeled with AF488-WGA (green). P*hfs*_CC_-*hfsH_HB_*, *C. crescentus* Δ*hfsH* cross*-*complemented with HfsH from *H. baltica* under the control of the *hfs* promoter; P*hfs*_HB_-*hfsH_CC_*, *H. baltica* Δ*hfsH cross-*complemented with HfsH from *C. crescentus* under the control of the *hfs* promoter. Exponential planktonic cultures were used to quantify the percentage of predivisional cells with holdfast. Data is expressed as the mean of three independent biological replicates with four technical replicates each. Error bars represent the standard error of the mean. **D.** Images showing merged phase and fluorescence channels of *C. crescentus* and *H. baltica* strains bound to glass slides. Holdfast is labeled with AF488-WGA (green). Exponential cultures were incubated on the glass slides for 1 h, unbound cells were washed off, and AF488-WGA was added to label bound holdfast. **E.** Merged phase and fluorescence channels of *H. baltica* Δ*hfsH* and *C. crescentus* Δ*hfsH* strains bound to glass slides. Holdfast is labeled with AF488-WGA (green). Exponential cultures were incubated on the glass slides for 1 h, unbound cells were washed off, and AF488-WGA was added to label bound holdfast. Strains carry native or cross-complemented HfsH under the control of the xylose inducible promoter for overexpression.

To test whether the cross-complemented strains restored biofilm formation, we quantified biofilm formed after 12 h (Fig. 5B). When we examined cross-complemented strains with HfsH under the control of the native promoter P*hfs*, we observed that P*hfs_CC_-hfsH_HB_* restored biofilm formation to WT levels in *C. crescentus* Δ*hfsH* (Fig. 5B). In contrast, P*hfs_HB_-hfsH_CC_* restored biofilm formation to only 20 % of WT levels in *H. baltica ΔhfsH* (Fig. 5B). When *hfsH_HB_* was overexpressed in *C. crescentus* from the P*xyl_CC_* promoter (*C. crescentus ΔhfsH* P*xyl_CC_-hfsH_HB_*), biofilm formation was increased to 150% of WT levels (Fig. 5B), similar to what has been observed with overexpression of HfsH_CC_ in *C. crescentus.*^27^ Overexpression of HfsH_CC_ using P*xyl_HB_* in *H. baltica* Δ*hfsH* (*H. baltica ΔhfsH* P*hfs_HB_-hfsH_CC_*) restored biofilm to WT levels (Fig. 5B). These results suggest that HfsH_HB_ may have higher levels of enzymatic activity than HfsH_CC_.

To analyze how cross-complementation of HfsH affects holdfast cohesion and anchoring, we labelled holdfasts from planktonic cultures with AF488-WGA. *C. crescentus ΔhfsH* P*hfs_CC_*-*hfsH_HB_* holdfasts were labelled with AF488-WGA and formed rosettes similar to the WT (95% of WT, Fig. 5C, left panels). In *H. baltica ΔhfsH* P*hfs_HB_-hfsH_CC_* we observed that approximately half of the stalked cells were labelled with AF488-WGA, however this labelling was weaker than the WT (50% of WT, Fig. 5C, right panels). These results indicate that P*hfs_CC_*-*hfsH_HB_* fully cross*-*complements *C. crescentus* Δ*hfsH*, while P*hfs_HB_*-*hfsH_CC_* only partially cross-complements *H. baltica ΔhfsH.* These results are in agreement with our observations for biofilm formation for these strains (Fig. 5B).

Since only half of *H. baltica* Δ*hfsH* P*hfs_HB_*-*hfsH_CC_* cells had faint AF488-WGA labelling and surface binding properties were not fully restored to WT levels (Fig. 5C, right panels), we hypothesized that holdfasts produced by this mutant may have been shed into the medium. Therefore, we tested whether the holdfast produced by this strain can bind to a glass surface by incubating cells on a coverslip at room temperature for 1 h. After incubation, unattached cells were washed off and AF488-WGA was added to label any holdfast bound to the coverslip. As expected, *C. crescentus ΔhfsH* cross-complemented with P*hfs_CC_*-*hfsH_HB_* produced holdfasts that bound to the coverslip and anchored the cells to the surface, like WT (Fig. 5D, left panels). We observed that holdfasts produced by *H. baltica ΔhfsH* P*hfs_HB_*-*hfsH_CC_* were not able to anchor the cells to the glass surface, although these holdfasts were able to bind to the coverslip (Fig. 5D, right panels). These results imply that expression of HfsH_CC_ from the *H. baltica hfsH* promoter was sufficient for restoration of holdfast surface binding by *H. baltica*, but insufficient to maintain interactions with the cell body. In addition, overexpression of either HfsH_CC_ or HfsH_HB_ in *C. crescentus* Δ*hfsH* and *H. baltica* Δ*hfsH* using P*xyl* restored holdfast binding properties to WT levels (Fig. 5E). These results suggest either that HfsH_HB_ and HfsH_CC_ have different levels of enzymatic activity or that their ability to deacetylate *H. baltica* holdfast is different, which could be contributing to the observed differences in *C. crescentus* and *H. baltica* holdfast binding properties.

### Increased HfsH expression improves binding in high ionic strength environments

In order to test whether HfsH plays an important role in holdfast binding at high ionic strength, we quantified binding of holdfasts purified from *C. crescentus* overexpressing HfsH_CC_ using the xylose promoter (P*xyl_CC_-hfsH_CC_*). To obtain cell-free holdfast samples and study holdfast without the contribution of the cell body, a holdfast-shedding mutation, Δ*hfaB*, was used as shown on Fig. 6A.^29,33^ Holdfasts from *C. crescentus* Δ*hfaB* Δ*hfsH* P*xyl_CC_-hfsH_CC_* (Fig. 6B, green line) tolerated higher ionic strengths than holdfasts purified from the control *C. crescentus* Δ*hfaB* strain (Fig. 6B, blue dashed line). These results suggest that increased expression of HfsH in *C. crescentus* improves ionic strength tolerance. When we overexpressed HfsH_HB_ in *C. crescentus* Δ*hfaB* Δ*hfsH*, we observed an increase in ionic strength tolerance similar to overexpression of HfsH_CC_, but not to the level observed in *H. baltica* (Fig. 6B, red, green and yellow lines). Our results suggest that holdfast polysaccharide deacetylation plays an important role in improving holdfast ionic strength tolerance, but it is not the only factor as increased HfsH_CC_ or HfsH_HB_ expression did not elevate *C. crescentus* holdfast binding to the level of *H. baltica*.

**Figure 6:**
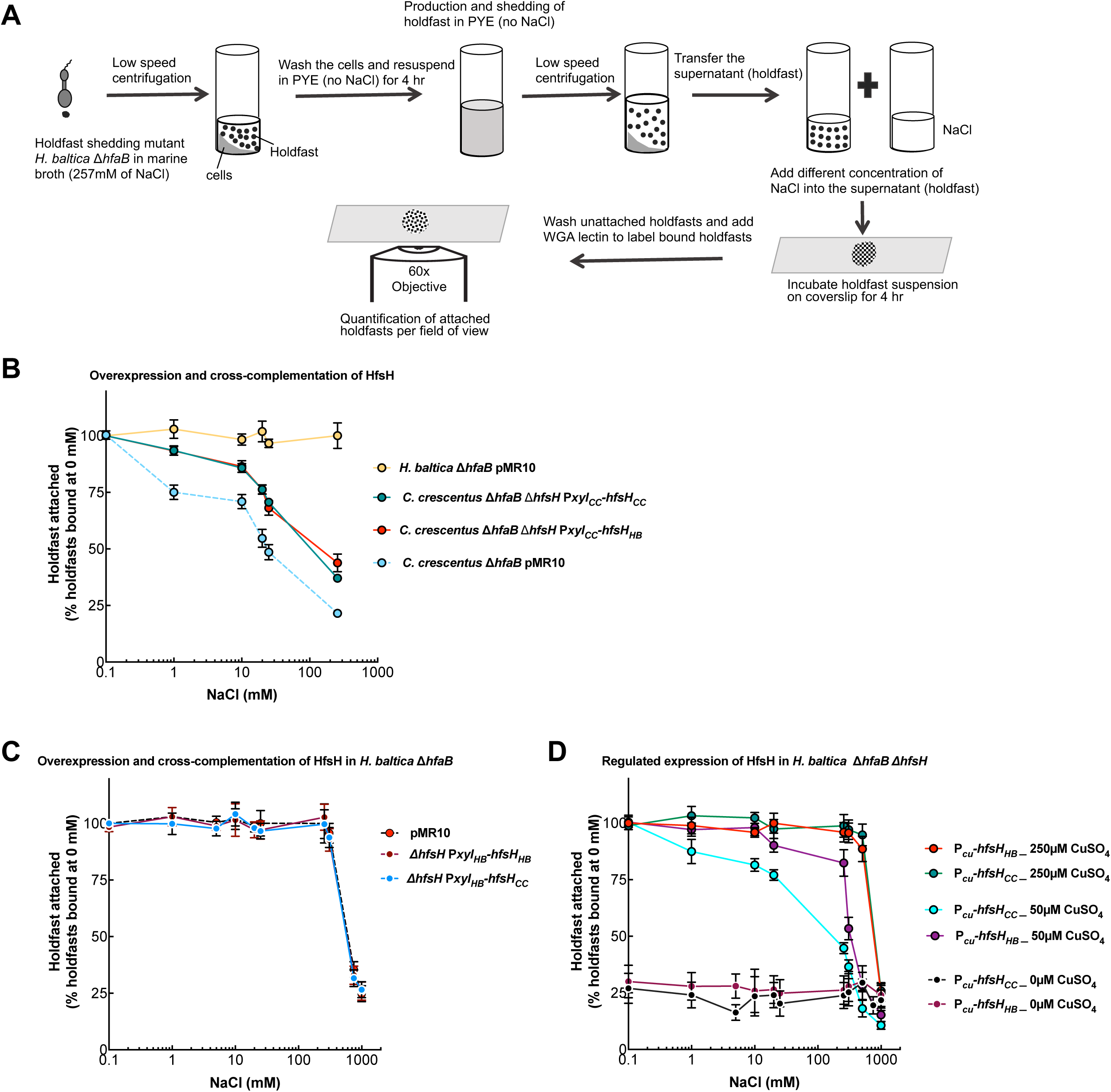
Increased holdfast deacetylation increases holdfast binding in high ionic strength environments. **A.** Schematic of experimental setup**. (B-D).** Cells were grown exponentially for 2 h in PYE with 0.03% xylose (A-B), or 0 µM, 50 µM, or 250 µM CuSO_4_ (C) and shed holdfast were collected from the culture supernatant. The purified holdfasts were mixed with different concentration of NaCl and incubated on glass slides for 4 h. The percentage of holdfasts bound per field of view were quantified at different concentrations of NaCl. The number of holdfasts bound per field of view at 0 mM NaCl was standardized to 100%. Data are expressed as an average of five independent biological replicates with five technical replicates each. Error bars represent the standard error of the mean. **B.** Purified holdfasts from *C. crescentus* Δ*hfaB* with pMR10 (empty vector, blue dashed line) as a control, *C. crescentus* Δ*hfaB* Δ*hfsH* complemented with HfsH from *C. crescentus* under the control of the xylose-inducible promoter (P*xyl*_CC_-*hfsH_CC_*, green), *C. crescentus* Δ*hfaB* Δ*hfsH* cross-complemented with HfsH from *H. baltica* under the control of the xylose-inducible promoter (P*xyl*_CC_-*hfsH_HB_*, red), and *H. baltica* Δ*hfaB* with pMR10 (empty vector, yellow). **C.** Purified holdfasts from *H. baltica* Δ*hfaB* with pMR10 (empty vector, black dashed line) as a control, *H. baltica* Δ*hfaB* Δ*hfsH* complemented with HfsH_HB_ under the control of the xylose-inducible promoter (P*xyl*_HB_-*hfsH_HB_*, maroon), and *H. baltica* Δ*hfaB* Δ*hfsH* cross*-*complemented with HfsH_CC_ under the control of the xylose-inducible promoter (P*xyl*_HB_-*hfsH_CC_*, blue). **D.** Purified holdfasts from *H. baltica* Δ*hfaB* Δ*hfsH* complemented with HfsH_HB_ under the control of the copper inducible promoter (P*_Cu_*-*hfsH_HB_*) and *H. baltica* Δ*hfaB* Δ*hfsH* cross*-*complemented with HfsH_CC_ under the control of the copper inducible promoter (P*_Cu_*-*hfsH_CC_*). P*_Cu_*-*hfsH_CC_* was induced with 0 µM (black), 50 µM (blue), and 250 µM CuSO_4_ (green), and P*_Cu_*-*hfsH_HB_* was induced with 0 µM (maroon), 50 µM (purple), and 250 µM CuSO_4_ (red).

We next examined the effect of cross-complementation with HfsH_CC_ in *H. baltica.* We had observed that expression of HfsH_CC_ in *H. baltica ΔhfsH* from P*hfs_HB_* failed to restore holdfast binding, but its overexpression using P*xyl* restored surface binding to the level observed in the WT (Fig. 5A-B). Therefore, we tested how holdfasts purified from *H. baltica* Δ*hfaB* overexpressing HfsH_CC_ responded to ionic strength. *H. baltica* Δ*hfaB* Δ*hfsH* P*xyl_HB_-hfsH_CC_* produced holdfasts that bound to glass slides to the same degree as holdfasts produced by control *H. baltica* Δ*hfaB* expressing regular levels of HfsH_HB_ (Fig. 6C, black dashed line and blue line). We also quantified holdfast binding from *H. baltica ΔhfaB* overexpressing HfsH_HB_ and did not observe a further increase in ionic strength tolerance (Fig. 6C, red line). These results suggest that *H. baltica* holdfasts either: (1) have maximized binding at native levels of HfsH_HB_ expression, or (2) have maximized holdfast deacetylation and further increases in HfsH_HB_ expression have no observable effects on holdfast binding.

We hypothesized that if increasing the level of HfsH expression increases *C. crescentus* holdfast binding at high ionic strength, then reducing the level of HfsH expression in *H. baltica* holdfast could reduce the ionic strength tolerance. To test this, we used the copper inducible promoter P*_Cu_* to control the expression of HfsH in *H. baltica* Δ*hfaB* Δ*hfsH.* We observed few holdfasts bound to the glass slide when HfsH_CC_ or HfsH_HB_ expression was not induced (Fig. 6D, maroon and black lines). However, at the highest level of induction of HfsH_HB_ and HfsH_CC_ (250 µM CuSO_4_), we observed full restoration of ionic strength tolerance (Fig. 6D, red and green lines). Next, we analyzed holdfast binding at an intermediate level of induction (50 µM CuSO_4_) because at this level of expression, we had observed 50% biofilm restoration and restoration of holdfast structure (Fig. 4B and S5A). Using Western blot analysis, we compared the level of expression of HfsH_HB_ and HfsH_CC_ at 50 µM CuSO_4_ and observed that they were expressed at similar levels (Fig. S5B). At intermediate levels of induction of HfsH_HB_, we observed a decrease in holdfast ionic strength tolerance compared to induction at 250 µM CuSO_4_ (Fig. 6D, purple curve). We observed a further decrease in holdfast ionic strength tolerance when HfsH_CC_ was induced with 50 µM CuSO_4_ compared to HfsH_HB_ (Fig. 6D, blue and purple curves). The effect of reducing HfsH expression was larger at high ionic strength than at low ionic strength, suggesting that holdfast polysaccharide deacetylation may play an important role in promoting holdfast binding at high ionic strength (Fig. 6D). These results suggest that HfsH_HB_ likely deacetylates *H. baltica* holdfast more efficiently than HfsH_CC_ and that marine *Caulobacterales* have optimized HfsH to augment holdfast binding at high ionic strength.

### HfsH is required for retention of holdfast thiols and galactose monosaccharides

In addition to polysaccharides, the *H. baltica* holdfast contain free thiol groups, suggesting that it contains proteins.^25^ Holdfast thiols require the presence of holdfast polysaccharides for cell association, as deletion of the glycosyltransferases essential for holdfast polysaccharide synthesis leads to loss of both holdfast polysaccharides and thiols.^25^ In addition to GlcNAc, *H. baltica* holdfasts contain galactose monosaccharides.^25^ In order to gain insights into how HfsH modifies holdfast properties, we analyzed its impact on these holdfast components.

In order to test whether holdfast thiols are present in the deacetylase mutant *H. baltica* Δ*hfsH*, we co-labeled exponential-phase cells with both AF488-WGA (green, GlcNAc) and AF594 conjugated to maleimide (AF594-Mal), which reacts with free thiols molecules (red). As expected, the WT cells were labeled with both AF488-WGA and AF594-Mal (Fig. 7A, left panels), indicating the presence of both holdfast polysaccharides and thiols. In contrast, the deacetylase mutant *H. baltica* Δ*hfsH* was not labelled with either AF488-WGA or AF594-Mal (Fig. 7A, right panels). We then varied the level of HfsH expression using *H. baltica* Δ*hfsH* P*_Cu_*-*hfsH*. Addition of CuSO_4_ to exponentially growing *H. baltica* WT cells with empty vector had no effect on labeling of holdfast polysaccharides (Fig. 7B, left panels). At the lowest level of induction (50 µM CuSO_4_), *H. baltica* Δ*hfsH* P*_Cu_-hfsH* holdfasts were labeled only by AF488-WGA while AF594-Mal failed to label holdfasts (Fig. 7B, right panels). We observed restored AF594-Mal labeling at the highest level of HfsH induction (250 µM CuSO_4_; Fig. 7B, right panels). These results indicate that HfsH expression is required for thiols contained molecules to associate with GlcNAc polysaccharides in the holdfast.

**Figure 7:**
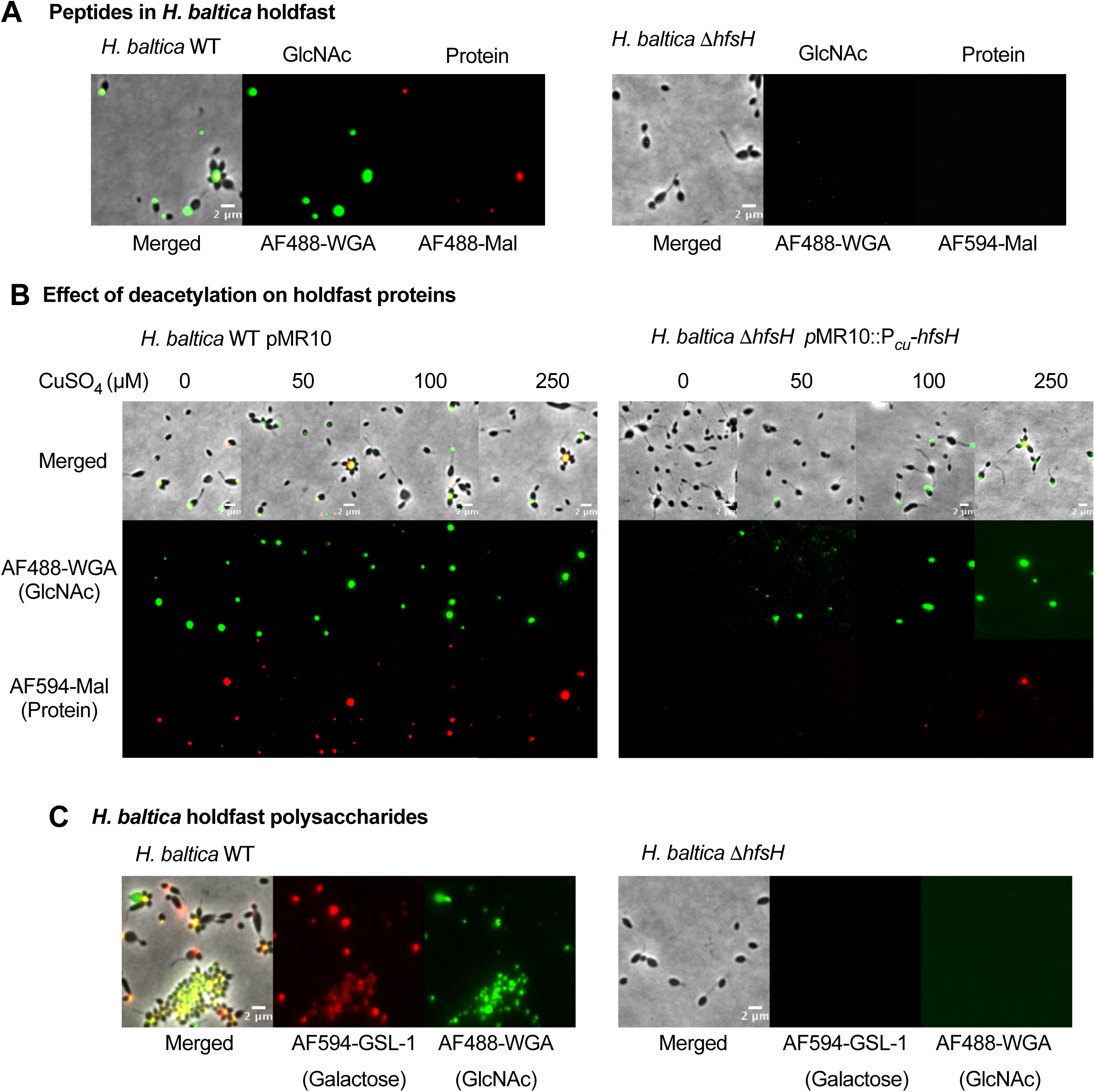
HfsH expression is required for interaction of holdfast thiols and galactose monosaccharides with cells. **A.** Representative images showing merged phase and fluorescence channels of *H. baltica* and *H. baltica* Δ*hfsH* holdfasts co-labeled with AF488-WGA (green, GlcNAc) to label polysaccharides and AF594-mal (red) to label free thiols. **B.** Representative images of *H. baltica* WT *and H. baltica* Δ*hfsH* complemented with pMR10::P*_Cu_*-*hfsH*. Holdfasts were co-labeled with AF488-WGA (green) and AF594-Mal (red). Exponential cultures were induced for 2 h with concentrations of CuSO_4_ ranging from 0 – 250 µM. **C.** Representative images showing merged phase and fluorescence channels of *H. baltica* and *H. baltica* Δ*hfsH* holdfasts co-labeled with AF488-WGA (green) and AF594-GSL-1 (red) to label GlcNAc and galactose in holdfast, respectively.

In order to test the effect of *hfsH* deletion on retention of the galactose component of holdfast, we co-labeled the cells with AF488-WGA (specific to GlcNAc, green) and AF594-conjugated *Griffonia simplicifolia* lectin 1 (AF594-GSL-1, specific to galactose, red). *H. baltica* WT holdfast was labelled with both AF488-WGA and AF594-GSL-1, but *H. baltica ΔhfsH* was not labeled with either AF594-GSL-1 or AF488-WGA (Fig. 7C). These results suggest that HfsH is also crucial for the retention of holdfast galactose components within the holdfast of *H. baltica*.

## DISCUSSION

Bacterial adhesion is influenced by many variables, including adhesin composition, surface properties, and environmental factors such as pH, temperature, fluid shear, and ionic strength.^10,23, 34–36^ *C. crescentus,* a freshwater *Caulobacterales*, produces holdfasts that are sensitive to ionic strength, while holdfasts from the marine *H. baltica* tolerate a higher ionic strength.^23,25^ In this study, we examined the influence of specific enzymatic modifications on the differing holdfast properties of these species. Specifically, we described the contributions of the polysaccharide deacetylase HfsH to holdfast binding at high ionic strength by comparing holdfasts from the freshwater *C. crescentus* and the marine *H. baltica.* The degree of deacetylation modifies holdfast polysaccharide physicochemical properties by introducing a partially positive charge on the resultant amine group, which is important for holdfast cohesive and adhesive properties.^27^

Our results showed that HfsH is important for biofilm formation and holdfast binding properties in *H. baltica*. We found that the *H. baltica* Δ*hfsH* mutant does not form rosettes or biofilms, and produces holdfasts that are impaired in surface binding and have a thread-like appearance, in contrast to the *C. crescentus* Δ*hfsH* mutant, which sheds small holdfasts that are capable of binding to glass surfaces. These observations suggest that there are differences in the role for holdfast deacetylation in *H. baltica versus C. crescentus.* These results also indicate that HfsH plays an important role in maintaining holdfast cohesive and adhesive properties in both species. It has been shown that polymers such as xylan and lignocellulose interact with surfaces using hydrogen bonds generated by deacetylation and the degree of deacetylation affects their interactions with polar surfaces.^37^ The degree of deacetylation of other polysaccharides such as chitin/chitosan, a polymer of GlcNAc, is important in altering their physicochemical properties. For example, deacetylation of chitin to generate chitosan increases its pK_a_ from 6.46 to 7.32.^30^ This pK_a_ change is due to an increase in free primary amines that are exposed by deacetylation. We believe that the polysaccharide deacetylase HfsH performs a similar function in modifying the holdfast polysaccharide.

Overexpression of HfsH_CC_ in *C. crescentus* increases holdfast adhesion without an increase in the size of holdfast.^27^ This suggests that *C. crescentus* holdfast polysaccharides are partially deacetylated and overexpression of HfsH_CC_ further increases the degree of deacetylation, in turn enhancing adhesion. In *H. baltica,* overexpression of HfsH_HB_ or HfsH_CC_ did not increase surface adhesion compared to WT. One interpretation of these results is that *H. baltica* holdfast polysaccharides are fully deacetylated by native levels of HfsH expression, and thus overexpression of HfsH has no additional effect. Alternatively, *H. baltica* holdfast binding is already maximized for out test conditions and thus an increase in deacetylation has no further positive effect. We hypothesized that if *H. baltica* has maximized its holdfast polysaccharide deacetylation, expression of HfsH below native levels would lead to a reduction in holdfast ionic strength tolerance. We showed that the level of HfsH expression correlates with holdfast binding in *H. baltica,* suggesting that *H. baltica* is exploiting deacetylation to optimize its binding in high ionic strength environments. Our results showed that HfsH_HB_ performs this function better than HfsH_CC,_ suggesting that there could be differences in HfsH_HB_ and HfsH_CC_ enzymatic activities. *H. baltica* produces larger holdfasts than *C. crescentus* and they contain additional sugars such as galactose,^25^ therefore an alternative hypothesis is that *C. crescentus* HfsH_CC_ might be less efficient at deacetylating *H. baltica* holdfast due to structural and compositional differences.

Interestingly, increasing expression of HfsH in *C. crescentus* leads to increased binding at high ionic strength. Cross-complementing *C. crescentus* Δ*hfsH* with overexpressed HfsH_HB_ produced holdfasts that had similar levels of increased ionic strength tolerance as those produced by overexpressed HfsH_CC_ as compared to WT, suggesting that HfsH_HB_ is capable of deacetylating *C. crescentus* holdfast polysaccharides. However, increased expression of HfsH_HB_ did not increase *C. crescentus* holdfast binding at high ionic strength to the level of *H. baltica,* implying that other factors also contribute to ionic strength tolerance. When we reduced the expression of HfsH_HB_ in *H. baltica,* we observed a decrease in ionic strength tolerance. A further decrease was observed when HfsH_CC_ was expressed at the same intermediate level in *H. baltica* compared to HfsH_HB_. These results suggest that, for *H. baltica* holdfast to overcome high ionic strength, there is a minimum level of deacetylation of holdfast polysaccharides that must be attained.

How holdfast interacts with surfaces remains unclear, but an electrostatic mechanism has been suggested.^23,25^ *C. crescentus* holdfast binding is affected by pH and NaCl.^23^ The mechanism by which NaCl disrupts electrostatic interactions between holdfast components and glass surfaces is unclear. High ionic strength has been shown to reduce the radius of the electrostatic force on a surface, which would lower the likelihood that holdfast polysaccharides are able to interact with the surface.^16^ It is also known that increasing ionic strength has no effect on holdfast that are already attached to a surface,^25^ suggesting that high ionic strength only impairs the initial interactions between the holdfast and the surface before a permanent bond is established. In *Pseudomonas putida*, it has been shown that high ionic strength alters the conformation of extracellular biopolymers.^38,39^ The polymer brush layer remains extended at low ionic strength, but upon an increase in ionic strength the brush layer becomes compacted, leading to an increase in the charge to mass ratio.^38,39^ This increase in charge to mass ratio ensures that the polysaccharides retain their electrostatic properties. This phenomenon could explain the need for a higher level of deacetylation of holdfasts from marine species compared to those of freshwater species, as deacetylation increases the proportion of charges on holdfast polysaccharides, as required for surface interactions at high ionic strength.

We conclude that degree of holdfast polysaccharide deacetylation is important in holdfast binding at high ionic strengths, and that the marine *Caulobacterales* have optimized deacetylation to overcome holdfast binding challenges in these environments. Generally, it seems like the degree of holdfast deacetylation and the degree of DOPA incorporation into the Mfps are equivalent strategies to adapt to increased ionic strength. We showed that the *H. baltica* Δ*hfsH* mutant lacks both galactose and thiol molecules, suggesting that these constituents require deacetylated holdfast to associate with the cell. Therefore, deacetylated GlcNAc, sugars other than GlcNAc, and/or putative thiol containing proteins might play a role in improving holdfast binding at high ionic strength. Validation of the presence of putative holdfast-associated proteins and their identification in *C. crescentus* and *H. baltica* will enable a better understanding of their role in holdfast binding in these environments.

## EXPERIMENTAL PROCEDURES

### Bacterial strains and growth conditions

The bacterial strains used in this study are listed in Table S1. *H. baltica* strains were grown in marine medium (Difco^TM^ Marine Broth/Agar reference 2216) except when studying the effect of ionic strength on holdfast binding, where they were grown in Peptone Yeast Extract (PYE) medium^19^ supplemented with 0 or 1.5% NaCl. *C. crescentus* was grown in PYE medium. Both *H. baltica* and *C. crescentus* strains were grown at 30 °C. When appropriate, kanamycin was added at 5 µg/ml to liquid medium and 20 µg/ml in agar plates. *H. baltica* strains with the copper inducible promoter were grown in marine broth supplemented with 0-250 µM of CuSO_4_. *H. baltica* strains with the xylose promoter were grown in marine broth supplemented with 0.03% xylose, while *C. crescentus* strains with the xylose promoter were grown in PYE broth supplemented with 0.03% xylose. *E. coli* strains were grown in lysogeny broth (LB) at 37 °C supplemented with 30 µg/ml of kanamycin in liquid medium or 25 µg/ml in agar plates, as appropriate.

### Strain construction

All the plasmids and primers used in this study are listed in Table S1 and S2, respectively. In-frame deletion mutants were obtained by double homologous recombination as previously described^40^ using suicide plasmids transformed into the *H. baltica* host strains by electroporation^41^ followed by sacB sucrose selection. Briefly, genomic DNA was used as the template to PCR-amplify 500 bp fragments immediately upstream and downstream of the gene to be deleted. The primers used for amplification were designed with 25 bp overlapping segments for isothermal assembly^42^ using the New England Biolabs NEBuilder tools for ligation into plasmid pNPTS139, which was digested using EcoRV-HF endonuclease from New England Biolabs. pNPTS139-based constructs were transformed into *α-*select *E. coli* for screening and sequence confirmation before introduction into the host *C. crescentus* or *H. baltica* strains by electroporation. Introduction of the desired mutation onto the *C. crescentus* or *H. baltica* genome was verified by sequencing.

For gene complementation, the pMR10 plasmid was cut with EcoRV-HF and 500 bp upstream of the gene of interest containing the promoter, as well as the gene itself, were designed using New England Biolabs NEBuilder tools and fragments were amplified and ligated into plasmid pMR10 as described above. The pMR10-based constructs were transformed into *α-*select *E. coli* for screening and sequence confirmation before introduction into the host *C. crescentus* or *H. baltica* strains by electroporation.

### Holdfast labeling using fluorescent lectins

Holdfast labeling with AF488 conjugated lectins (Molecular Probes) was performed as previously described^25^ with the following modifications. Overnight cultures were diluted in fresh medium to an OD_600_ of 0.2 and incubated for 4 h to an OD_600_ of 0.6 – 0.8. AF488 conjugated lectins were added to 100 µl of the exponential culture to a final concentration of 0.5 µg/ml and incubated at room temperature for 5 min. 5 µl of the labeled culture was spotted onto a glass cover slide, overlaid with a 1.5 % (w/v) agarose (Sigma-Aldrich) pad in water, and visualized by epifluorescence microscopy. Imaging was performed using an inverted Nikon Ti-E microscope with a Plan Apo 60X objective, a GFP/DsRed filter cube, an Andor iXon3 DU885 EM CCD camera, and Nikon NIS Elements imaging software with 200 ms exposure time. Images were processed in ImageJ.^43^

### Short-term adherence and biofilm assays

This assay was performed as previously described^25^ with the following modifications. For short-term binding assays, exponential cultures (OD_600_ of 0.6 - 0.8) were diluted to an OD_600_ of 0.4 in fresh marine broth, added to 24-well plates (1 ml per well), and incubated with shaking (100 rpm) at room temperature for 4 h. For biofilm assays, overnight cultures were diluted to an OD_600_ of 0.1, added to a 24-well plate (1 ml per well), and incubated at room temperature for 12 hours with shaking (100 rpm). In both set-ups, OD_600_ was measured before the wells were rinsed with distilled H_2_O to remove non-adherent bacteria, stained using 0.1% crystal violet (CV), and rinsed again with dH_2_O to remove excess CV. The CV was dissolved with 10% (v/v) acetic acid and quantified by measuring the absorbance at 600 nm (A_600_). Biofilm formation was normalized to A_600_ / OD_600_ and expressed as a percentage of WT.

### HfsH expression using a copper inducible promoter

Strains bearing copper inducible plasmids were inoculated from freshly grown colonies into 5 ml marine broth containing 5 μg/ml kanamycin and incubated with shaking (200 rpm) at 30°C overnight. Overnight cultures were diluted in fresh marine broth to OD_600_ of 0.1 and incubated until an OD_600_ of 0.4 was reached, where copper sulfate dissolved in marine broth was added to a final concentration of 0-250 µM. For holdfast labeling, AF488 conjugated lectins were added to 100 µl of exponential culture to a final concentration of 0.5 µg/ml and incubated at room temperature for 5 min. 5 µl of the labeled culture was spotted on glass cover slide, overlaid with a 1.5 % (w/v) agarose (Sigma-Aldrich) pad in water, and visualized by epifluorescence microscopy. Imaging was performed using an inverted Nikon Ti-E microscope with a Plan Apo 60X objective, a GFP/DsRed filter cube, an Andor iXon3 DU885 EM CCD camera and Nikon NIS Elements imaging software with 200 ms exposure time. Images were processed in ImageJ.^43^

For short term binding and biofilm assays, the induced cultures and controls (OD_600_ = 0.4) were incubated with shaking (100 rpm) at room temperature for 4 - 12 h. Then, OD_600_ was measured before the wells were rinsed with distilled H_2_O to remove non-adherent bacteria, stained using 0.1% crystal violet (CV), and rinsed again with dH_2_O to remove excess CV. The CV was dissolved with 10% (v/v) acetic acid and quantified by measuring the absorbance at 600 nm (A_600_). Biofilm formation was normalized to A_600_ / OD_600_ and expressed as a percentage of WT.

### Visualization of holdfasts attached on a glass surface

Visualization of holdfast binding to glass surfaces was performed as described previously^25^ with the following modifications. *H. baltica and C. crescentus* strains grown to exponential phase (OD_600_ = 0.4 – 0.6) were incubated on washed glass coverslips at room temperature in a saturated humidity chamber for 4 - 8 h. After incubation, the slides were rinsed with dH_2_O to remove unbound cells, holdfasts were labelled using 50 µl of fluorescent AF488/594 conjugated lectins at a concentration of 0.5 µg/ml, and cover slides were incubated at room temperature for 5 min. Then, excess lectin was washed off and the cover slide was topped with a glass coverslip. Holdfasts were imaged by epifluorescence microscopy using an inverted Nikon Ti-E microscope with a Plan Apo 60X objective, a GFP/DsRed filter cube, an Andor iXon3 DU885 EM CCD camera and Nikon NIS Elements imaging software with 200 ms exposure time. Images were processed in ImageJ.^43^

### Holdfast synthesis by time-lapse microscopy on soft agarose pads

*H. baltica* holdfast synthesis was observed in live cells on agarose pads by time-lapse microscopy as described previously^25^ with some modifications. A 1 µl aliquot of exponential-phase cells (OD_600_ of 0.4 – 0.8) induced with 0 – 250 µM CuSO_4_ was placed on top of a 0.8% agarose pad in marine broth with 0.5 µg/ml of AF488-WGA. The pad was overlaid with a coverslip and sealed with VALAP (Vaseline, lanolin and paraffin wax). Time-lapse microscopy images were taken every 5 min for 12 h using an inverted Nikon Ti-E microscope and a Plan Apo 60X objective, a GFP/DsRed filter cube, and an Andor iXon3 DU885 EM CCD camera. Time-lapse movies were processed using ImageJ.^43^

### Holdfast synthesis in a microfluidic device by time-lapse microscopy

This experiment was performed as previously described^25^ with the following modifications. Cell cultures were grown to mid-exponential phase (OD_600_ = 0.4-0.6) and induced with 0 – 250 µM CuSO_4_. Then, 200 µl of culture was diluted into 800 µl of fresh marine broth with 0 – 250 µM CuSO_4_ in the presence of 0.5 µg/ml AF488-WGA for holdfast labeling. One ml of the cell culture was then flushed into a microfluidic device containing a 10 µm high linear chamber fabricated in PDMS (Polydimethylsiloxane) as described previously.^44^ After injection of the cells into the microfluidic chamber, the flow rate was adjusted so that attachment could be observed under static conditions or low flow rate of 1.4 µl/min.

Time-lapse microscopy was performed using an inverted Nikon Ti-E microscope and a Plan Apo 60X objective, a GFP/DsRed filter cube, an Andor iXon3 DU885 EM CCD camera, and Nikon NIS Elements imaging software. Time-lapse videos were collected over a period of 5.5 h at 20-second intervals. Cell attachment was detected at the glass-liquid interface within the microfluidic chamber using phase contrast microscopy, while holdfast synthesis was detected using fluorescence microscopy. Time-lapse movies were processed using ImageJ.^43^

### Holdfast labeling using fluorescently labeled maleimide and lectin

Alexa Flour conjugated Maleimide C_5_ (AF488-mal, ThermoFisher Scientific) and AF594-WGA (Molecular Probes) were both added to 100 µl of exponential culture to a final concentration of 0.5 µg/ml and incubated at room temperature for 5 min. 5 µl of the labeled culture was spotted onto a glass cover slide, overlaid with a 1.5 % (w/v) agarose (Sigma-Aldrich) pad in water, and visualized by epifluorescence microscopy. Imaging was performed using an inverted Nikon Ti-E microscope with a Plan Apo 60X objective, a GFP/DsRed filter cube, an Andor iXon3 DU885 EM CCD camera, and Nikon NIS Elements imaging software with 200 ms exposure time. Images were processed in ImageJ.^43^

### Effect of ionic strength on holdfast binding

Visualization of attachment of purified holdfasts to surfaces at different ionic strengths was performed as described previously^25^ with the following modifications. Briefly, exponential cultures of strains carrying *hfsH* under the control of the copper inducible promoter (P*_Cu_*), xylose inducible promoters (P*xyl*), or controls (P*hfs*) were grown to late exponential phase (OD_600_ = 0.6 – 0.8) in PYE with 1.5% (w/v) NaCl for *H. baltica* strains, or PYE with no NaCl for *C. crescentus* strains with 0 – 250 µM CuSO_4_ or 0.03% xylose. The cells were collected by centrifugation for 30 min at 4,000 x *g* and resuspended in PYE with 0 – 250 µM CuSO_4_ or 0.03% xylose and incubated for 2 h at 30 °C. Then, the cells were again collected by centrifugation as above and 100 µl of the resultant supernatant, containing holdfasts shed by the cells, were mixed with 100 µl of NaCl in PYE to a final concentration of 0 - 1000 mM of NaCl. 50 µl of the mixture was incubated on washed glass coverslips at room temperature in a saturated humidity chamber for 4 - 12 h. After incubation, the slides were rinsed with dH_2_O to remove unbound material and holdfast were visualized with AF conjugated lectins (Molecular Probes). Imaging was performed using an inverted Nikon Ti-E microscope with a Plan Apo 60X objective, a GFP/DsRed filter cube, an Andor iXon3 DU885 EM CCD camera, and Nikon NIS Elements imaging software with 200 ms exposure time. Images were processed in ImageJ.^43^ The number of holdfasts bound per field of view was quantified using MicrobeJ.^45^

### Western blot analysis

Cell lysates were prepared from exponentially growing cultures (OD_600_ = 0.6-0.8) as previously described^27^ with the following modifications. The equivalent of 1.0 ml of culture at an OD_600_ of 0.6-0.8 was centrifuged at 16 000 × *g* for 5 min at 4 °C. The supernatant was removed, and cell pellets were resuspended in 50 µl of 10mM Tris pH 8.0, followed by the addition of 50 µl of 2x SDS sample buffer. Samples were boiled for 5 min at 100 °C before being run on a 12% (w/v) polyacrylamide gel and transferred to a nitrocellulose membrane. Membranes were blocked for 30 min in 5% (w/v) non-fat dry milk in TBST (20 mM Tris, pH 8, 0.05% (w/v) Tween 20), and incubated at 4 °C overnight with primary antibodies. Anti-FLAG tag and McpA antibodies were used at a concentration of 1:10 000. Then, a 1:10 000 dilution of secondary antibody, HRP-conjugated goat anti-rabbit immunoglobulin, was incubated with the membranes at room temperature for 2 h. Membranes were developed with SuperSignal West Dura Substrate (Thermo Scientific, Rockford, IL).

## ACKNOWLEDGEMENTS

We thank the members of the Brun laboratory for the discussion and providing critical comments on the manuscript. This study was supported by grant R35GM122556 from the National Institutes of Health to YVB. YVB is supported by a Canada 150 Research Chair in Bacterial Cell Biology from the Canadian Institutes of Health Research.

## AUTHOR CONTRIBUTION

YVB and NKC designed the research. NKC performed the research. YVB and NKC analyzed the data. YVB and NKC wrote the paper.

## Supporting Information

### The HfsK acetyltransferase is not required for holdfast adhesion and biofilm formation in *H. baltica*

*C. crescentus* HfsK is involved in holdfast modification, although its role is unclear.^1^ *C*. *crescentus* Δ*hfsK* produces holdfasts that are less adhesive, are not cohesive, and are shed into the medium.^1^ In a glass surface binding assay, *C*. *crescentus* Δ*hfsK* produces holdfasts that adhere to glass, but fails to anchor cells in place^1^ To test whether *hfsK* and its paralogs play a role in biofilm formation in *H. baltica,* we generated in-frame deletion mutants of *H. baltica hfsK* and its paralogs *hbal_1607* and *hbal_1184*. The *H. baltica ΔhfsK* mutant showed no defect in biofilm formation after 12 h incubation at room temperature (Fig. S1A). We observed similar results for the *H. baltica* Δ*hbal_1607* and the *H. baltica* Δ*hbal_1184* mutants, as well as the triple deletion mutant *H. baltica ΔhfsK Δhbal_1607 Δhbal_1184* (Fig. S1A). These results indicate that HfsK and its paralogs are not involved *H. baltica* biofilm formation, in contrast to what has been reported for *C. crescentus*^1^ and *H. baltica hfsH* mutant (Fig. S1A).

As holdfast is required for biofilm formation in *C. crescentus* and *H. baltica,*^2–4^ we probed for the presence of holdfasts using fluorescent Alexa Fluor 488 (AF488) conjugated <u>w</u>heat <u>g</u>erm <u>a</u>gglutinin (WGA) lectin that specifically binds to the GlcNAc component of the holdfast polysaccharide.^4^ In exponentially growing planktonic cultures, *C. crescentus* WT cells produced holdfasts that bound AF488-WGA and formed cell-cell aggregates mediated by holdfasts, called rosettes (Fig. S1B, left panel). *C. crescentus ΔhfsK* produced holdfasts which variably were associated with the cell or were shed into the medium (Fig. S1B, left panel, white and blue arrows), as previously shown.^1^ Deletion of *hfsK* in *H. baltica* had no effect on AF488-WGA binding to holdfast, rosette formation, or holdfast shedding (Fig. S1B), consistent with its lack of an effect on biofilm formation (Fig. S1A).

In order to test whether *H. baltica* HfsK is involved in holdfast anchoring, we spotted exponentially growing cultures on a glass coverslip and incubated for 1 h at room temperature to allow for binding to the coverslip. Unbound cells were removed by washing, and AF488-WGA was added to label holdfasts that remained attached to the coverslip. As a control, *C. crescentus* and *H. baltica* WT cells were incubated with coverslips, and adherent holdfasts were labeled with AF488-WGA (Fig. S1C). *C. crescentus ΔhfsK* holdfasts were bound to coverslips but appeared to be spread over the surface, covering a greater area than WT and suggesting that they may be less cohesive (Fig. S1C), in agreement with previous studies.^1^ These holdfasts also failed to anchor *C. crescentus ΔhfsK* cells to the surface (3% of WT, Fig. S1C). In comparison, mutants with deletion of *hfsK* and its paralogs in *H. baltica* produced holdfasts that were bound to the glass surface and formed rosettes similarly to WT (Fig. S1C right panel). Interestingly, deletion of the *H. baltica hfsK* paralog *hbal*_*1184* led to the generation of large cellular aggregates that formed independently of holdfast biogenesis (Fig. S1D). These cells had morphological defects and were surrounded by debris that may have resulted from cell lysis, indicating that Hbal_1184 is likely involved in a different polysaccharide biosynthetic pathway that contributes to cellular viability. We conclude that HfsK and its paralogs do not contribute to *H. baltica* holdfast binding properties under our assay conditions (Fig. S1A-C).

## EXPERIMENTAL PROCEDURES

### Bacterial strains and growth conditions

The bacterial strains used in this study are listed in Table S1. *H. baltica* strains were grown in marine medium (Difco^TM^ Marine Broth/Agar reference 2216) except when studying the effect of ionic strength on holdfast binding, where they were grown in Peptone Yeast Extract (PYE) medium^5^ supplemented with 0 or 1.5% NaCl. *C. crescentus* was grown in PYE medium. Both *H. baltica* and *C. crescentus* strains were grown at 30 °C. When appropriate, kanamycin was added at 5 µg/ml to liquid medium and 20 µg/ml in agar plates. *H. baltica* strains with the copper inducible promoter were grown in marine broth supplemented with 0-250 µM of CuSO_4_. *H. baltica* strains with the xylose promoter were grown in marine broth supplemented with 0.03% xylose, while *C. crescentus* strains with the xylose promoter were grown in PYE broth supplemented with 0.03% xylose. *E. coli* strains were grown in lysogeny broth (LB) at 37 °C supplemented with 30 µg/ml of kanamycin in liquid medium or 25 µg/ml in agar plates, as appropriate.

### Strain construction

All the plasmids and primers used in this study are listed in Table S1 and S2, respectively. In-frame deletion mutants were obtained by double homologous recombination as previously described^6^ using suicide plasmids transformed into the *H. baltica* host strains by electroporation^7^ followed by sacB sucrose selection. Briefly, genomic DNA was used as the template to PCR-amplify 500 bp fragments immediately upstream and downstream of the gene to be deleted. The primers used for amplification were designed with 25 bp overlapping segments for isothermal assembly^8^ using the New England Biolabs NEBuilder tools for ligation into plasmid pNPTS139, which was digested using EcoRV-HF endonuclease from New England Biolabs. pNPTS139-based constructs were transformed into *α-*select *E. coli* for screening and sequence confirmation before introduction into the host *C. crescentus* or *H. baltica* strains by electroporation. Introduction of the desired mutation onto the *C. crescentus* or *H. baltica* genome was verified by sequencing.

### Holdfast labeling using fluorescent lectins

Holdfast labeling with AF488 conjugated lectins (Molecular Probes) was performed as previously described^2^ with the following modifications. Overnight cultures were diluted in fresh medium to an OD_600_ of 0.2 and incubated for 4 h to an OD_600_ of 0.6 – 0.8. AF488 conjugated lectins were added to 100 µl of the exponential culture to a final concentration of 0.5 µg/ml and incubated at room temperature for 5 min. 5 µl of the labeled culture was spotted onto a glass cover slide, overlaid with a 1.5 % (w/v) agarose (Sigma-Aldrich) pad in water, and visualized by epifluorescence microscopy. Imaging was performed using an inverted Nikon Ti-E microscope with a Plan Apo 60X objective, a GFP/DsRed filter cube, an Andor iXon3 DU885 EM CCD camera, and Nikon NIS Elements imaging software with 200 ms exposure time. Images were processed in ImageJ.^9^

### Short-term adherence and biofilm assays

This assay was performed as previously described^2^ with the following modifications. For short-term binding assays, exponential cultures (OD_600_ of 0.6 - 0.8) were diluted to an OD_600_ of 0.4 in fresh marine broth, added to 24-well plates (1 ml per well), and incubated with shaking (100 rpm) at room temperature for 4 h. For biofilm assays, overnight cultures were diluted to an OD_600_ of 0.1, added to a 24-well plate (1 ml per well), and incubated at room temperature for 12 hours with shaking (100 rpm). In both set-ups, OD_600_ was measured before the wells were rinsed with distilled H_2_O to remove non-adherent bacteria, stained using 0.1% crystal violet (CV), and rinsed again with dH_2_O to remove excess CV. The CV was dissolved with 10% (v/v) acetic acid and quantified by measuring the absorbance at 600 nm (A_600_). Biofilm formation was normalized to A_600_ / OD_600_ and expressed as a percentage of WT.

### HfsH expression using a copper inducible promoter

Strains bearing copper inducible plasmids were inoculated from freshly grown colonies into 5 ml marine broth containing 5 μg/ml kanamycin and incubated with shaking (200 rpm) at 30°C overnight. Overnight cultures were diluted in fresh marine broth to OD_600_ of 0.1 and incubated until an OD_600_ of 0.4 was reached, where copper sulfate dissolved in marine broth was added to a final concentration of 0-250 µM. For holdfast labeling, AF488 conjugated lectins were added to 100 µl of exponential culture to a final concentration of 0.5 µg/ml and incubated at room temperature for 5 min. 5 µl of the labeled culture was spotted on glass cover slide, overlaid with a 1.5 % (w/v) agarose (Sigma-Aldrich) pad in water, and visualized by epifluorescence microscopy. Imaging was performed using an inverted Nikon Ti-E microscope with a Plan Apo 60X objective, a GFP/DsRed filter cube, an Andor iXon3 DU885 EM CCD camera and Nikon NIS Elements imaging software with 200 ms exposure time. Images were processed in ImageJ.^9^

### Visualization of holdfasts attached on a glass surface

Visualization of holdfast binding to glass surfaces was performed as described previously^2^ with the following modifications. *H. baltica and C. crescentus* strains grown to exponential phase (OD_600_ = 0.4 – 0.6) were incubated on washed glass coverslips at room temperature in a saturated humidity chamber for 4 - 8 h. After incubation, the slides were rinsed with dH_2_O to remove unbound cells, holdfasts were labelled using 50 µl of fluorescent AF488/594 conjugated lectins at a concentration of 0.5 µg/ml, and cover slides were incubated at room temperature for 5 min. Then, excess lectin was washed off and the cover slide was topped with a glass coverslip. Holdfasts were imaged by epifluorescence microscopy using an inverted Nikon Ti-E microscope with a Plan Apo 60X objective, a GFP/DsRed filter cube, an Andor iXon3 DU885 EM CCD camera and Nikon NIS Elements imaging software with 200 ms exposure time. Images were processed in ImageJ.^9^

### Western blot analysis

Cell lysates were prepared from exponentially growing cultures (OD_600_ = 0.6-0.8) as previously described^10^ with the following modifications. The equivalent of 1.0 ml of culture at an OD_600_ of 0.6-0.8 was centrifuged at 16 000 × *g* for 5 min at 4 °C. The supernatant was removed, and cell pellets were resuspended in 50 µl of 10mM Tris pH 8.0, followed by the addition of 50 µl of 2x SDS sample buffer. Samples were boiled for 5 min at 100 °C before being run on a 12% (w/v) polyacrylamide gel and transferred to a nitrocellulose membrane. Membranes were blocked for 30 min in 5% (w/v) non-fat dry milk in TBST (20 mM Tris, pH 8, 0.05% (w/v) Tween 20), and incubated at 4 °C overnight with primary antibodies. Anti-FLAG tag and McpA antibodies were used at a concentration of 1:10 000. Then, a 1:10 000 dilution of secondary antibody, HRP-conjugated goat anti-rabbit immunoglobulin, was incubated with the membranes at room temperature for 2 h. Membranes were developed with SuperSignal West Dura Substrate (Thermo Scientific, Rockford, IL).

## SUPPORTING TABLES

**TABLE S1:**
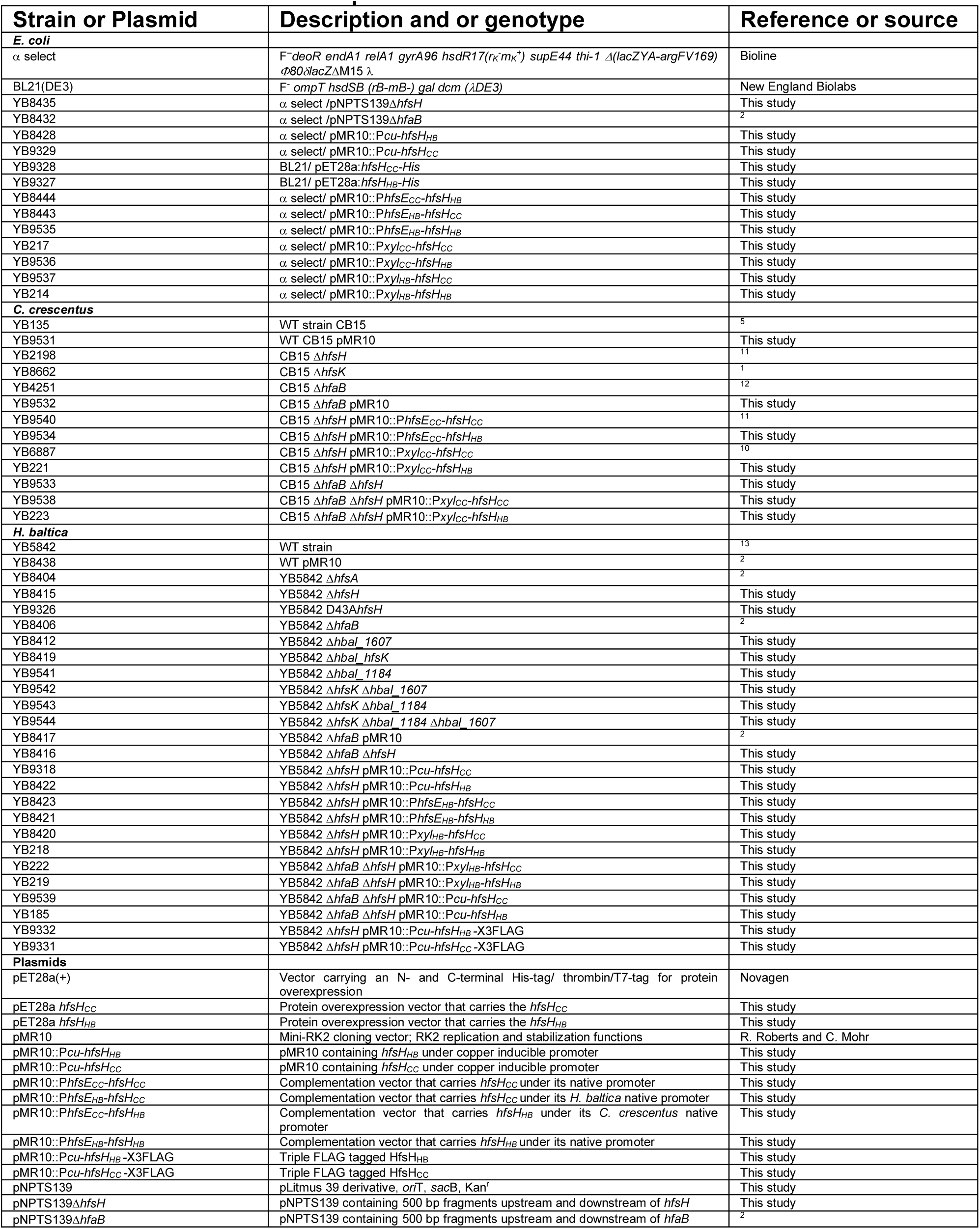
Bacterial strains and plasmids

**Tables S2:**
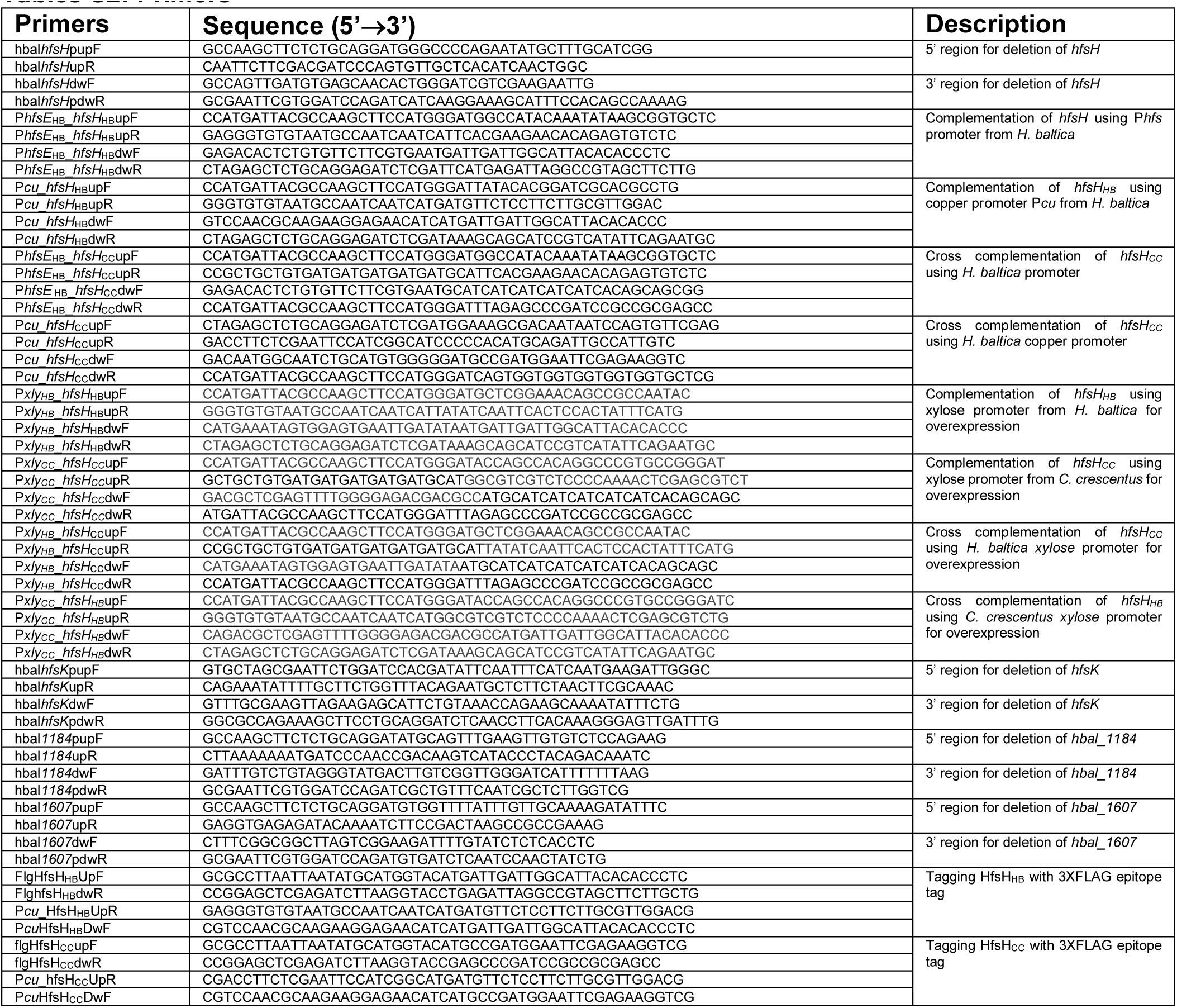
Primers

## SUPPORTING FIGURES AND FIGURE LEGENDS

**Figure S1:**
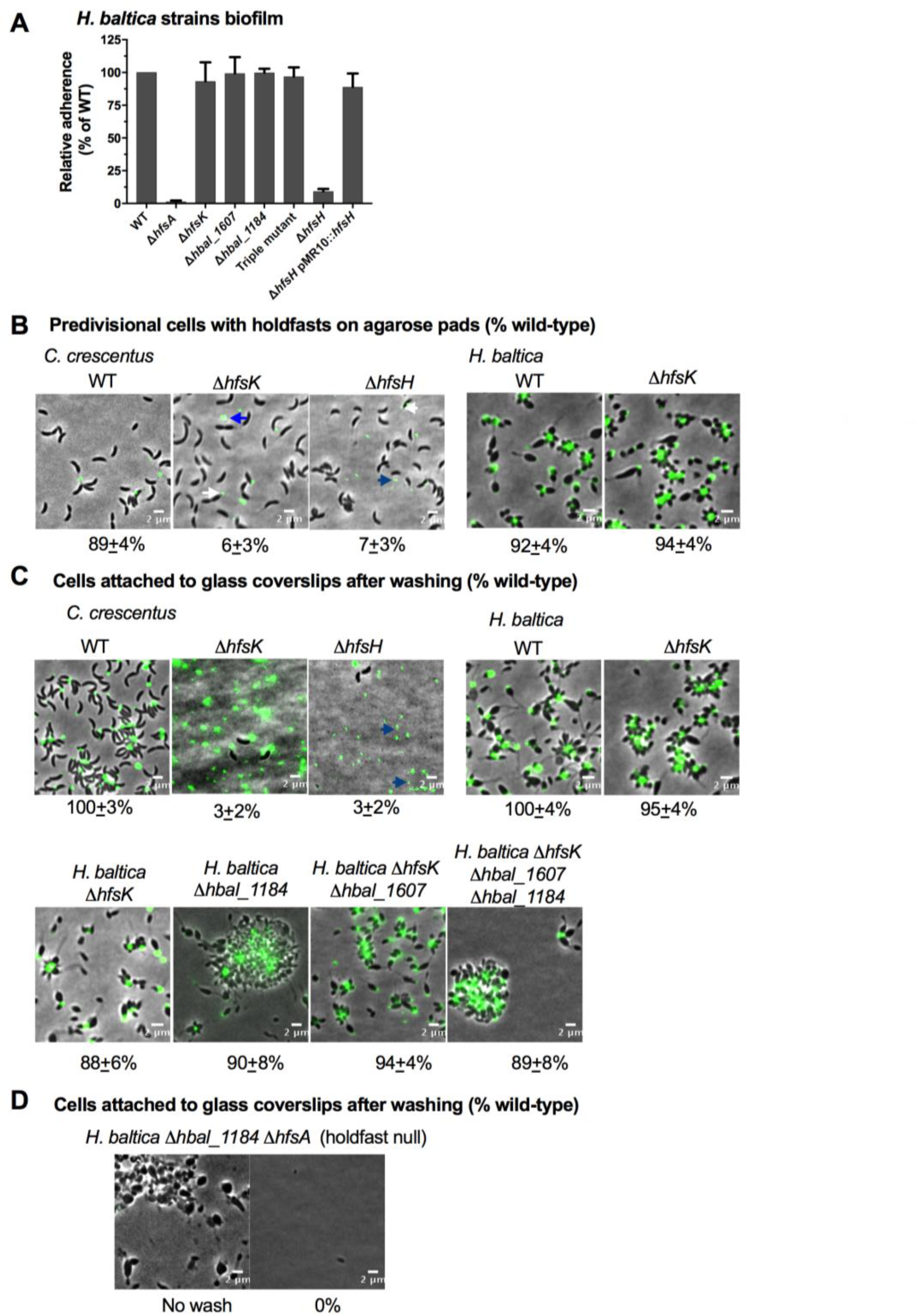
The HfsK acetyltransferase is not required for holdfast adhesion and biofilm formation in *H. baltica*. **A.** Quantification of biofilm formation by the crystal violet assay after incubation for 12 h, expressed as a mean percent of WT crystal violet staining. Holdfast null strain *ΔhfsA* was used as a negative control. Error is expressed as the standard error of the mean of three independent biological replicates with four technical replicates each. The triple mutant is *H. baltica ΔhfsK Δhbal_1607 Δhbal_1184.* **B.** Representative images showing merged phase and fluorescence channels of the indicated *C. crescentus* and *H. baltica* strains on agarose pads. Holdfast is labeled with AF488-WGA (green). White arrows indicate holdfasts attached to the Δ*hfsH* and Δ*hfsK* cells, and blue arrows indicate holdfast shed into the medium. Exponential planktonic cultures were used to quantify the percentage of predivisional cells with holdfast. Data are expressed as the mean of three independent biological replicates with four technical replicates each. Error bars represent the standard error of the mean. A total of 3,000 cells were quantified per replicate using MicrobeJ (Images for the WT and *hfsH* are from Fig. 2). **C-D.** Representative images showing merged phase and fluorescence channels of *C. crescentus* and *H. baltica* strains bound to a glass coverslip. Exponential cultures were incubated on the glass slides for 1 h, washed to remove unbound cells, and holdfast were labelled with AF488-WGA (green). Blue arrows indicate surface-bound holdfasts shed by *hfsH* mutants. The data showing quantification of cells bound to the glass coverslip are the mean of two biological replicates with five technical replicates each. Error is expressed as the standard error of the mean using MicrobeJ (Images for the WT and *hfsH* are from Fig. 2).

**Figure S2:**
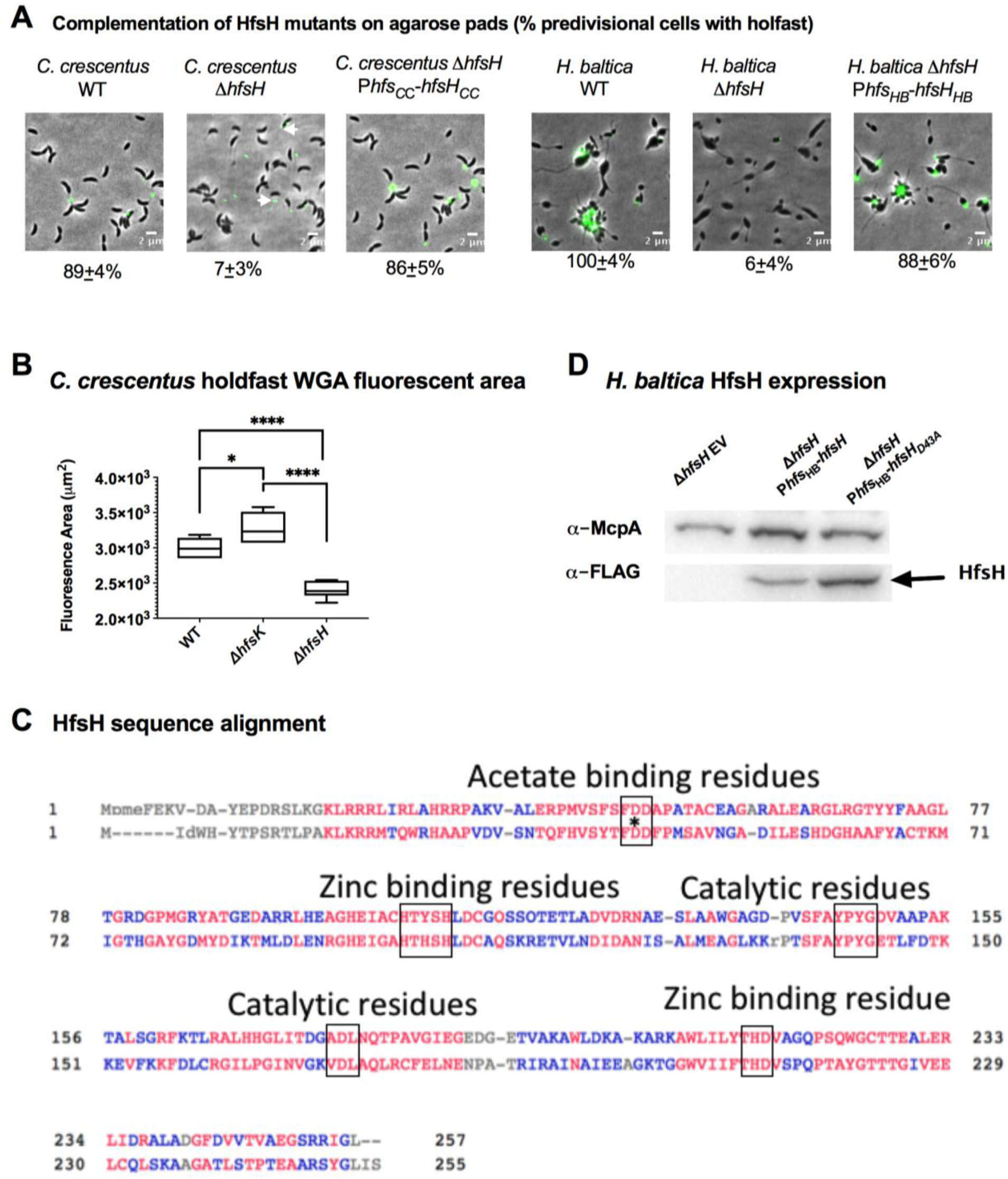
*H. baltica* and *C. crescentus* modification enzymes HfsK and HfsH. **A.** Images showing merged phase and fluorescence channels of the indicated strains of *C. crescentus* (left) and *H. baltica* (right). Holdfast is labeled with AF488-WGA (green). Exponential planktonic cultures were used to quantify the percentage of predivisional cells with holdfast. Data is expressed as the mean of three independent biological replicates with four technical replicates each. Error bars represent the standard error of the mean. **B.** Box plot showing the area of AF488-WGA fluorescence from holdfast produced by *C. crescentus* strains. Data is the mean of four biological replicates. The horizontal bar represents the median, the box represents 25^th^ and 75^th^ percentile, and the whiskers represent the full range of data. * and *** represent *P values* <0.1 and <0.0001 **C.** Alignment of the *C. crescentus* and *H. baltica* HfsH amino acid sequences with conserved carbohydrate esterase family 4 (CE4) motifs indicated by rectangles. The conserved residue involved in acetate binding is indicated with an asterisk (D48 in *C. crescentus* HfsH and D43 in *H. baltica* HfsH). **D.** Western blots of whole cell lysates of the indicated strains showing the expression level of FLAG-tag fusions of HfsH and HfsH_D43A_. McpA levels were monitored as a loading control.

**Figure S3:**
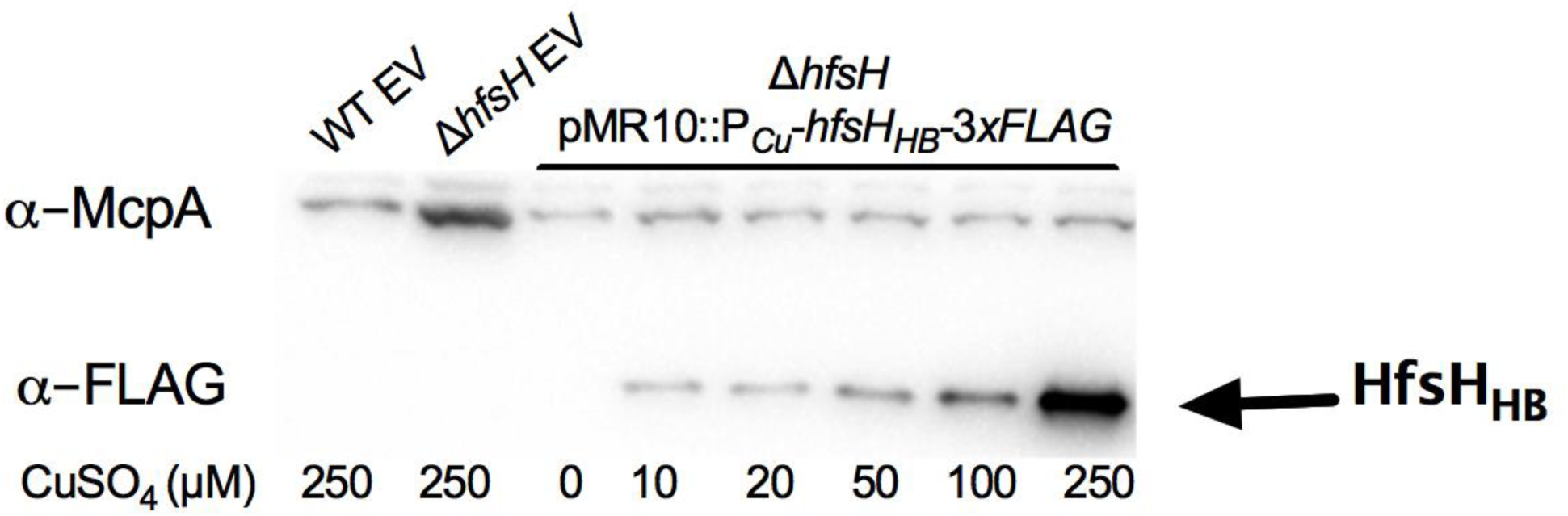
HfsH expression using a copper inducible promoter (P*_Cu_*) in *H. baltica*. Western blots of whole cell lysates showing the expression levels of FLAG-tagged HfsH under the control of the copper inducible promoter after 4 h of induction with 0 – 250 µM CuSO_4_. McpA levels were monitored as a loading control.

**Figure S4:**
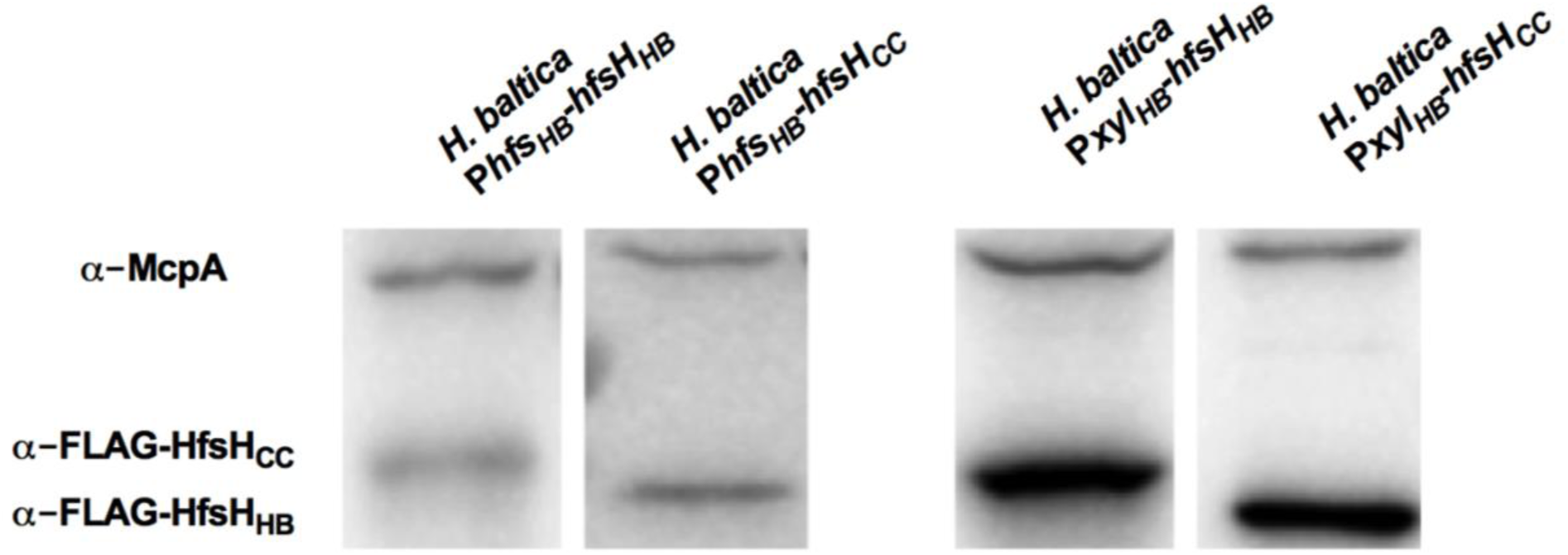
Levels of HfsH_HB_ and HfsH_CC_ expression using P*hfs* and P*xyl* promoters in *H. baltica*. Western blots of whole cell lysates showing the expression levels of FLAG-tagged HfsH_HB_ and HfsH_CC_ under the control of native holdfast synthesis (P*hfs)* and xylose-inducible (P*xyl*) promoters after 4 h of induction with 0.03% xylose. McpA levels were monitored as a loading control.

**Figure S5:**
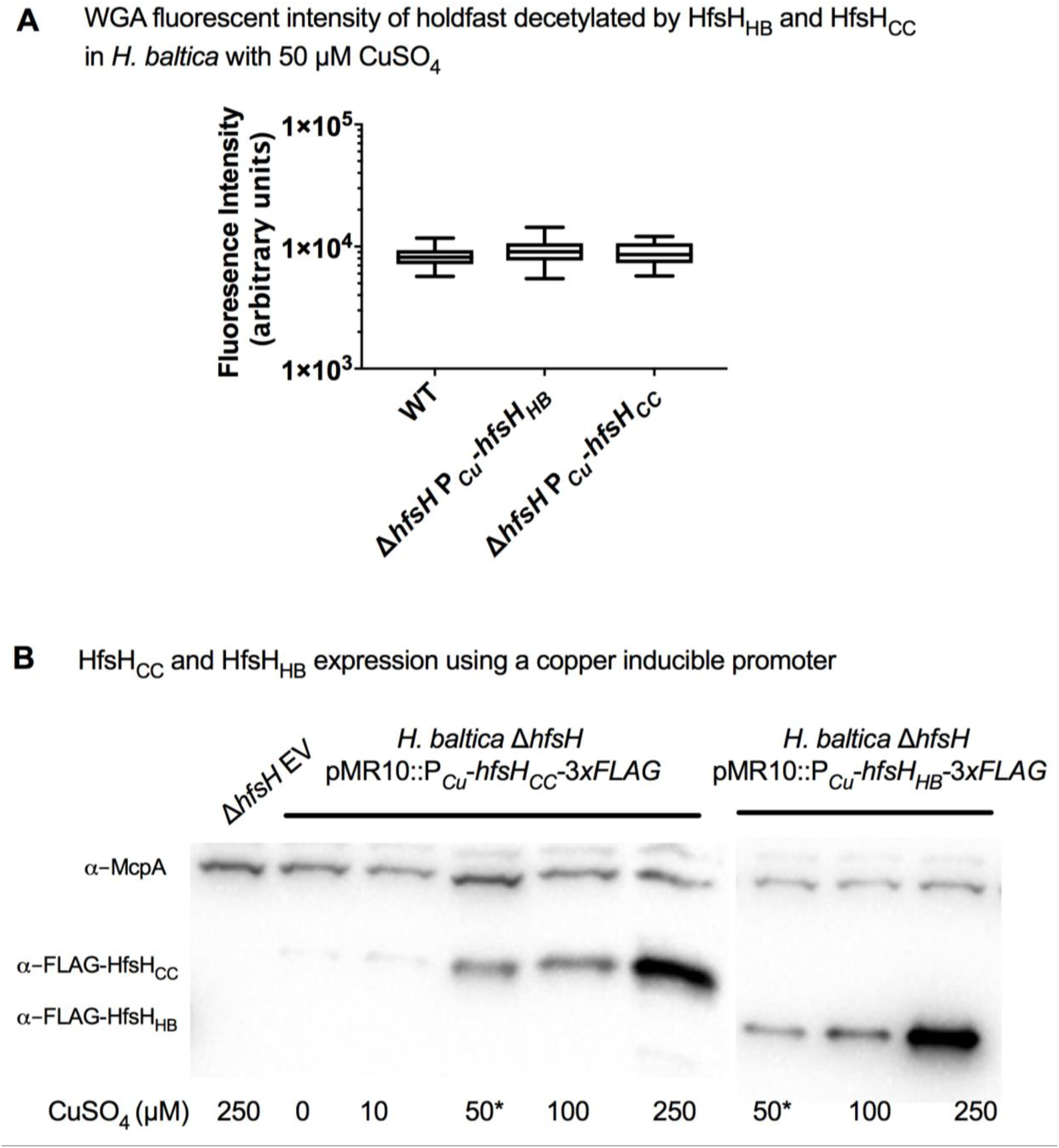
Cross-complementation of HfsH_HB_ and HfsH_CC_ using the copper inducible promoter. **A.** Box plot showing the integrated intensity of AF488-WGA fluorescence from holdfast produced by *H. baltica* Δ*hfsH* pMR10::P*_Cu_*-*hfsH_HB_* and cross-complemented *H. baltica* Δ*hfsH* pMR10::P*_Cu_*-*hfsH_CC_* at 50 µM CuSO_4_ induction for 4 h. Data is the mean of four biological replicates. The horizontal bar represents the median, the box represents 25^th^ and 75^th^ percentile, and the whiskers represent the full range of data. **B.** Western blots of whole cell lysates showing the expression level of FLAG-tagged HfsH_HB_ and HfsH_CC_ under the control of the copper inducible promoter after 4 h of induction with 0 – 250 µM CuSO_4_. McpA levels were monitored as a loading control. The star indicates HfsH induction at 50 µM CuSO_4_, used in comparing *H. baltica* and *C. crescentus* holdfast binding.

## SUPPORTING MOVIE LEGENDS

**Movie S1**

**A-B.** Time-lapse video of *H. baltica* WT and *H. baltica* Δ*hfsH* on soft agarose pads. Exponential cultures were placed on soft agarose pads containing holdfast-specific AF488-WGA (green) and covered with a glass coverslip. Images were collected every 5 min for 12 h. **C-D.** Time-lapse video of *H. baltica* WT and *H. baltica* Δ*hfsH* in microfluidic channels. Exponential cultures with holdfast-specific AF488-WGA (green) were injected into the microfluidic chambers and flow was turned off. Images were collected every 20 sec for 5.5 h.

**Movie S2**

**A-C.** Time-lapse videos of *H. baltica* Δ*hfsH* pMR10::P*_Cu_-hfsH* in microfluidic channels with holdfast labeled with AF488-WGA (green). Exponential cultures were induced with 0 µM, 50 µM, or 250 µM CuSO_4_, mixed with AF488-WGA, injected into the microfluidic chambers, and allowed to bind for 30 min. Thereafter, the flow rate was adjusted to 1.4 ul/min. Images were collected every 20 sec for 1 h.

